# A Comparative Study of Novel Microscopic Entities in Rock Samples: Implications for Panspermia and the Origin of Life

**DOI:** 10.1101/2025.05.07.652759

**Authors:** Juan Manuel González Castaño, Diana Melisa Montoya Alzate, Fredy Abelardo Zapata, Arcángel Montes Ortiz, Mauricio Hoyos, Ricardo Rangel-Martínez

## Abstract

This study presents the discovery and characterization of novel microscopic structures found on the surface of a rock sample collected from a stream in Pereira, Risaralda, Colombia. The structures, termed microcóndrulos, exhibit spherical morphologies atypical of known terrestrial entities. Qualitative and semi-quantitative chemical analyses were performed using scanning electron microscopy with energy-dispersive X-ray spectrometry (SEM-EDS). Results revealed a complex elemental composition, with up to forty-eight different elements detected, including C, O, N, Si, Ti, V, Ni, La, and Ce, among others. The microcóndrulos were differentiated from known terrestrial contaminants such as pollen, bacteria, and various protist groups. Comparative analysis with previously reported stratospheric samples and micrometeorites suggests a possible non-terrestrial origin. These findings contribute to the ongoing discussion on the diversity of life and the potential for alternative biogenesis. A taxonomic proposal for these new entities is presented for the first time.

## Introduction

The expectation of finding life on other worlds distracts us from the possibility of discovering new life forms on our own planet. Astrobiology is the scientific field that uses concepts and methods from biology, chemistry, geology, physics, planetary sciences, and astronomy to address fundamental aspects of life (Yamagishi, 2019). The emergence of astrobiology as a scientific term occurred in the late 1990s (1998), when the NASA Astrobiology Institute (NAI) began its work (Staley, 2003).

In this context, the main objective of astrobiology is the study of the origin, evolution, and distribution of life in the universe (Irvine, 2015), which makes it necessary to address issues such as the discovery of primitive or more evolved life forms and the understanding of their origin and diversity.

The origin of life on Earth, one of the main topics of astrobiology, is considered one of the possible explanations for the origin of life, along with the panspermia hypothesis. The latter theory assumes that life on Earth originated outside our planet and that its evolution is attributable to an interplanetary or interstellar transfer of “seeds” or “spores” (de Vera, 2015; Wesson, 2010; Hoyle and Wickramasinghe, 1981; Sergievskii, 2020).

There are several possibilities for how panspermia might occur. A first variant suggests that life in the universe has expanded due to a constant flow of rocks (lithopanaspermia). Other interpretations are based on the expansion of life through radiation pressure from the sun and other celestial bodies (radiopanspermia), the intervention of an advanced civilization and spacecraft (directed panspermia), and a natural process favored by certain circumstances (specifically, the existence of comets). (Wesson, 2010). Finally, it is pertinent to mention more recent variants such as necropanspermia, which is based on the transfer of inactivated viruses and bacteria throughout the universe; neopanspermia, which maintains that the arrival of life on Earth from outer space did not occur only in the past but continues to appear in the present; pathospermia, based on the arrival of diseases to Earth from outside; and archaeapanspermia, which suggests the possibility that dormant organisms on Earth arrived from space and returned to it (Wainwright et al., 2021).

Life is found in spaces and conditions where researchers previously thought it was impossible to exist. A number of previous studies have indicated that bacterial and fungal populations can be found in the stratosphere. Some have found these populations in the lower stratosphere (Griffin, 2004, 2008; Smith et al., 2010), while other studies have reported results from the upper stratosphere (Imshenetsky et al., 1978; Harris et al., 2002; Wainwright et al., 2003; Yang et al., 2008, 2009; Shivaji et al., 2009).

Wainwright et al. (2013a, 2013c, 2013d, 2013e) reported the isolation of some entities considered biological, which were collected in the stratosphere in 2013. These entities consist of biomorphs, most notably a sphere composed of titanium, vanadium, carbon, and nitrogen (the latter three elements were found in small quantities), which the authors hypothesized contained a biological protoplast (Wainwright et al., 2013g), a probable fragment of a diatom frustule (cell wall) (found at an altitude between 22 and 27 km), some filaments (Wainwright et al., 2013f), and other probable unidentified organisms (Wainwright et al., 2014). The same authors cited in this paragraph concluded that these entities and forms did not originate from Earth.

Likewise, Wainwright et al. (2013b), analyzing a sample from a meteorite that fell near the city of Polonnaruwa (Sri Lanka), found carbonaceous microspheres (fossilized microbial structures), including diatoms, embedded in the rock, ruling out the possibility that they were contaminants (Wickramasinghe et al., 2015). Wainwright et al. (2013h) similarly stated that fossilized worm-like bioforms were also found in that meteorite, concluding that they were likely bacteria. The findings of possible diatoms in this meteorite were reaffirmed years later (Wainwright and Wickramasinghe, 2021). Subsequently, Wainwright et al. (2015a) conducted two new experiments to collect samples in the stratosphere, through which they discovered new biological entities composed of carbon, oxygen, and other elements such as calcium, iron, magnesium, sulfur, and silicon.

Another important fact related to the topic addressed in this article is that, in the early 2000s, researchers from Cardiff University (United Kingdom) together with Indian researchers studied a particle found in the stratosphere, at an altitude of 41 km. Through various tests, they determined that the particle contained living matter, more specifically DNA, and that this genetic material corresponded to a bacterial organism (Wainwright, 2015).

Other researchers, such as Hoover et al. (2005), building on the initial work of Rossignol-Strick and Barghoorn (1971), studied a series of shapes, such as membranes and spiral-like structures, as well as spherical hollow microstructures showing well-defined walls (in acid macerated extract) present in a carbonaceous meteorite called Orgueil. Furthermore, the presence of numerous organic compounds in another meteorite called Murchison has also been reported (Cronin et al. 1988; Schmitt-Koplin et al. 2010).

Reciently, in an article published in the Journal of Modern Physics called: “Extraterrestrial Life in the Thermosphere: Plasmas, UAP, Pre-Life, Fourth State of Matter”, Joseph R. et al., investigates the presence and characteristics of “plasmas” in the Earth’s thermosphere, observed during NASA space shuttle missions. These self-illuminated entities, reaching up to a kilometer in size, exhibit behaviors reminiscent of multicellular organisms, including attraction to electromagnetic radiation, shape-shifting, and interactions with satellites. The study analyzes flight path trajectories, revealing varied velocities and directional changes, with some plasmas demonstrating what appears to be “hunter-predatory” behavior.

The analysis suggests these plasmas may represent a unique form of pre-life, potentially capable of incorporating elements common in space to synthesize RNA. Observed behaviors include congregating in the thermosphere, descending into thunderstorms, and interacting with satellites, generating electromagnetic activity. These phenomena might account for many Unidentified Aerial-Anomalous Phenomenon (UFO-UAP) sightings, as plasmas attracted to electromagnetic activity descend into the lower atmosphere.

This research posits that these plasmas constitute a fourth state of matter with life-like properties, distinct from biological entities but exhibiting complex behaviors and potential for pre-biological processes. The study proposes further investigation into the composition and behavior of these plasmas, suggesting locations for observation and capture to better understand their role in space and their potential connection to unexplained aerial phenomena.

Based on the information provided in the introduction, it is pertinent to question the real possibility of discovering new microscopic biological entities of possible extraterrestrial origin on our own planet. Astrobiology, as a scientific field, has broadened its focus to include not only the search for life on other worlds, but also the understanding of the limits and diversity of life on Earth.

The theory of panspermia, along with its various variants (lithopanspermia, radiopanspermia, necropanspermia, among others), suggests that life could have originated outside of Earth and arrived here through various mechanisms, such as the transport of “seeds” or “spores” in rocks or comets. The findings of biological entities in the stratosphere and in meteorites, composed of elements such as titanium, vanadium, carbon, and nitrogen, as well as the identification of possible bacteria and worm-like structures, support the idea that extraterrestrial life could be present on our planet.

In addition, research on plasmas in the thermosphere, observed during NASA space shuttle missions, raises the possibility of a pre-biotic life form capable of synthesizing RNA and exhibiting complex behaviors. These plasmas, attracted by electromagnetic activity, could descend into the lower atmosphere, which would explain some unidentified aerial phenomena.

Considering all these findings and theories, it is reasonable to consider the possibility that there are microscopic biological entities of extraterrestrial origin yet to be discovered on Earth, which invites us to investigate further and question our preconceptions about the limits of life.

The objective of this study is to present evidence to the scientific community of a new typology of microchondrule-like entities with spherical morphology present on the surface of a rock found in a stream in a Colombian municipality, which could be associated with a new class of biological entities not yet classified, with a possible non-terrestrial origin.

## Materials and Methods

### Description of Procedures and Analysis

The study material is a rock found in March 2020 in a stream called Quebrada San José, located in the area of the same name in the municipality of Pereira, Department of Risaralda, Colombia. Its origin is unknown, and no similar material exists in the area.

### Nomenclature

Figure 3 shows the statistical analysis: three groups or “clusters” (C) of microspheres classified as C1, C2, and C3. The statistical analysis was performed considering three aspects of the microspheres:

1. Chemical composition
2. Morphology
3. Sizes or diameters

A general characterization of the rock was performed based on its most distinctive features visible to the naked eye using zoom binocular magnifying glasses and a scanning electron microscope (SEM), as well as a laboratory analysis of the mineralogical composition using X-ray diffractometry (XRD).

For the detailed study, image analysis and semi-quantitative chemical analyses were performed using a scanning electron microscope (SEM) with energy-dispersive X-ray spectrometry (EDS). The sample was prepared under high and low vacuum (working pressure of 6·10-4 Pa).

Sixty-one independent microspheres were characterized at different sampling times. The morphology of potential new entities, similar to microchondrules or microspheres, was captured using a high-resolution energy-dispersive scanning microscope (SEM-EDS). The equipment used was an FEI Quanta 250 model with a tungsten electron source with a resolution of 3.0 nm at 15 kV and 30 kV in its different operating modes. Different sampling and observation sessions were conducted directly on the rock material, with 11 sessions.

A database was created with qualitative and semi-quantitative information from each of the samplings performed. The analysis used relative weight percentages (wt%) of the chemical elements detected by scanning electron microscopy. In some cases, it was possible to take samples of the microsphere on more than one occasion; therefore, the simple average of the weight percentages (wt%) found was used in the database. Each microsphere was classified with a code that included the letter of the general group (S: sphere) and a consecutive sample number. Qualitative information included the morphology and characteristics visible in the images captured by the scanning electron microscope (SEM), in which distinctive features could be differentiated using the external appearance as an individual classification (brain, granular, lobulated, or smooth).) and including their sphericity (spheroid or ovoid), the presence of protuberances (with or without), the presence of spots (with or without), the presence of scales (with or without) and the apparent size within the group (small < 5 µm, medium 5 µm < < 15 µm or large > 15 µm).

## Statistical Analysis

Due to the characteristics of the available data, the multivariate cluster analysis methodology was chosen, performed independently for both quantitative and qualitative data using the statistical software Infostat (Di Rienzo et al., 2019).

In the cluster analysis, unstandardized weight percentages (wt%) were used as the variables that performed best for performing the different groupings. The squared Euclidean distance was used as a measure. Hierarchical clustering was performed using Ward’s method or minimum variance method (Ward, 1963), and the non-hierarchical K-means method was used, taking as a reference the number of clusters suggested in the hierarchical method at 50% of the calculated distance.

For cluster analysis of qualitative data, the positive matching coefficient was used as the similarity coefficient, and the function for obtaining distance measures from similarity indices (S) was used as a complement to the absolute value of similarity (1 - abs (S)). Finally, Ward’s method was applied as a hierarchical clustering method.

X-ray diffraction

Chemical analysis

## Results and discussion

Macroscopic Description and Mineralogical Composition.

When wet, the rock’s veins and microveins change color from white to green (photo attached).

Some fusion surfaces can be observed on the front of the rock.

The texture of the rock is very strange from a geological point of view. The front (what I called the rock’s front) presents a large number of veins and microveins, while the back of the rock shows almost no veins or microveins.

Using a normal digital microscope at 1600x magnification, some very strange textures were observed, resembling very thin curved filaments, other very thin curvilinear filaments, and other straight black filaments. Other features included very small black circular inclusions and others that were whitish.

Small, very luminous rectangles emitting multicolored beams (could they be quasi-crystals) of a composition undetermined or microdiamonds.

Despite being a rock with a low weight under ambient conditions (151.3 g) and dimensions of 7.3 cm high, 5.9 cm wide, and 2.5 cm thick (Fig. 1), based on laboratory analyses, it was possible to establish that the rock had both an igneous and metamorphic composition: basaltic, composed mainly of olivine, hornblende, plagioclase (albite), and magnesite hornblende; and metamorphic, metabasalt (composed mainly of olivine, chorlomite (fayalite), actinolite, plagioclase, hornblende, and quartz); with a texture of very unusual greenish-gray veinlets composed of microcrystals of plagioclase (albite-labradorite) and microcrystals of hornblende and tourmaline adhered or bonded together by a micromaterial of organic composition (carbon (C), oxygen (O), silicon (Si), and aluminum (Al). The above analysis was performed taking into account the results obtained with X-ray diffractometry (XRD) (Figure 2).

**Figure 1:**
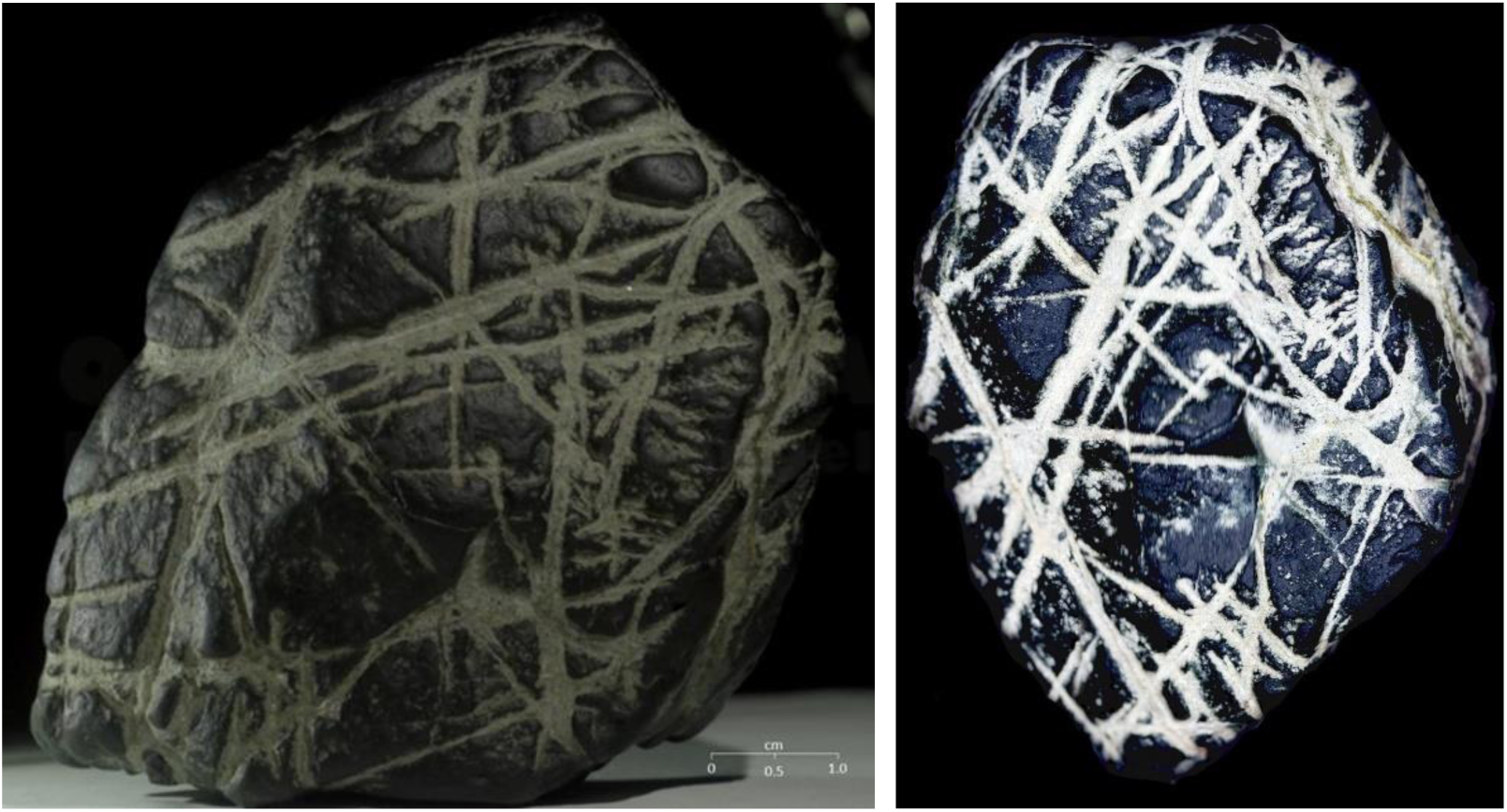
Rock with microchondrules.

**Fig. 2:**
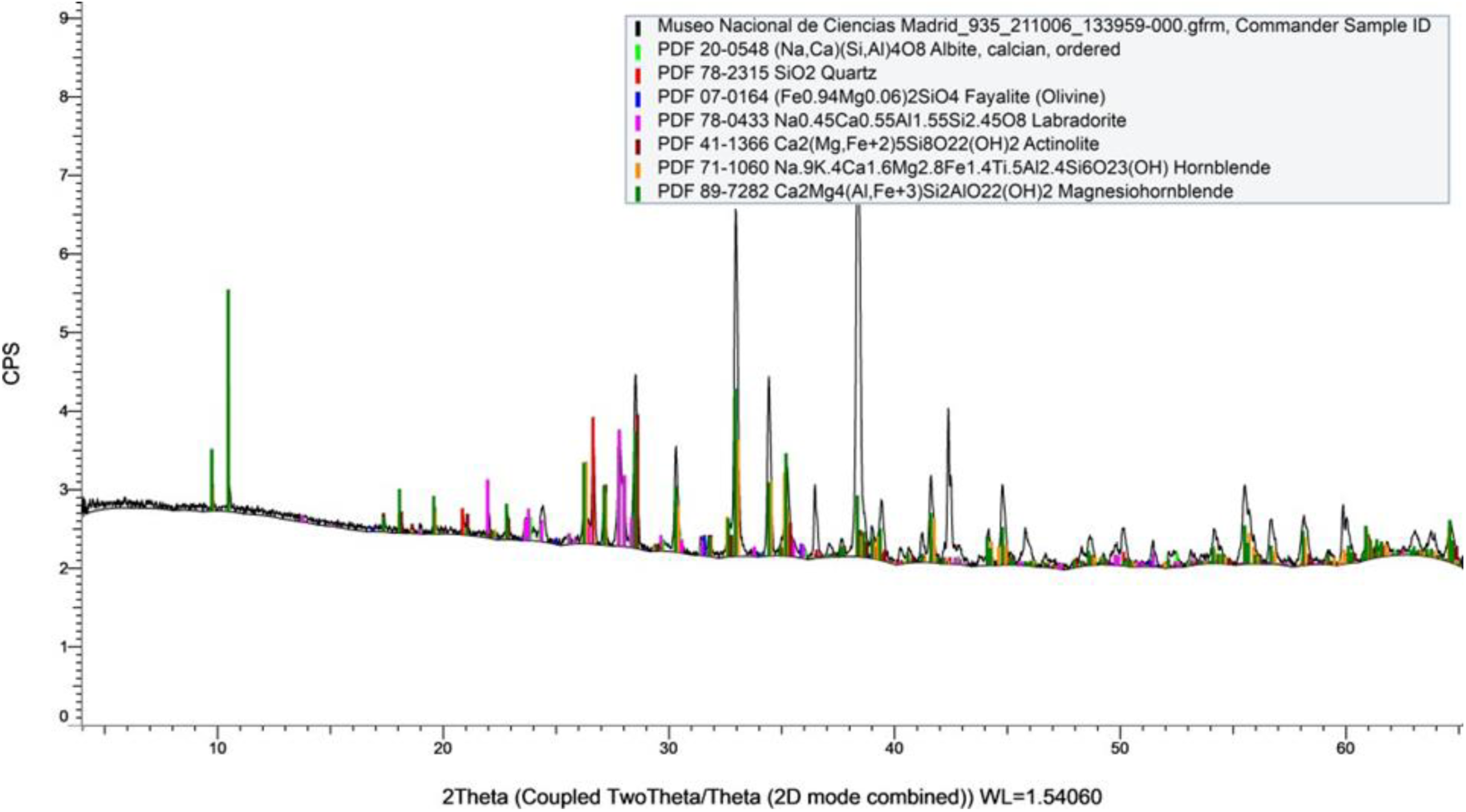
Analysis of the mineralogical composition of the rock with X-ray diffractometry.

### Groups according to semiquantitative chemical composition

A total of sixty-one independent microspheres were characterized at different sampling times. The analysis shows the possible grouping of the microspheres into three main classes (Fig. 3), which were determined by the elements present and their respective weight percentage (wt%) estimated in the microanalysis report. Of the clustering techniques used, it was decided to present a hierarchical analysis because both agreed on the classification. Based on the SEM results, forty-eight chemical elements from the periodic table were identified, with the presence of nonmetallic elements essential for life, such as carbon, oxygen, nitrogen, phosphorus, and sulfur; halogens, such as fluorine, chlorine, and bromine; and thirty-seven metallic elements belonging to the alkaline, alkaline earth, transition, and post-transition metal groups, most notably Al, Fe, Ca, Mg, Ni, Na, and Ga; and rare earth elements from the lanthanide group, such as Ce and La (Table 1).

**Fig. 3:**
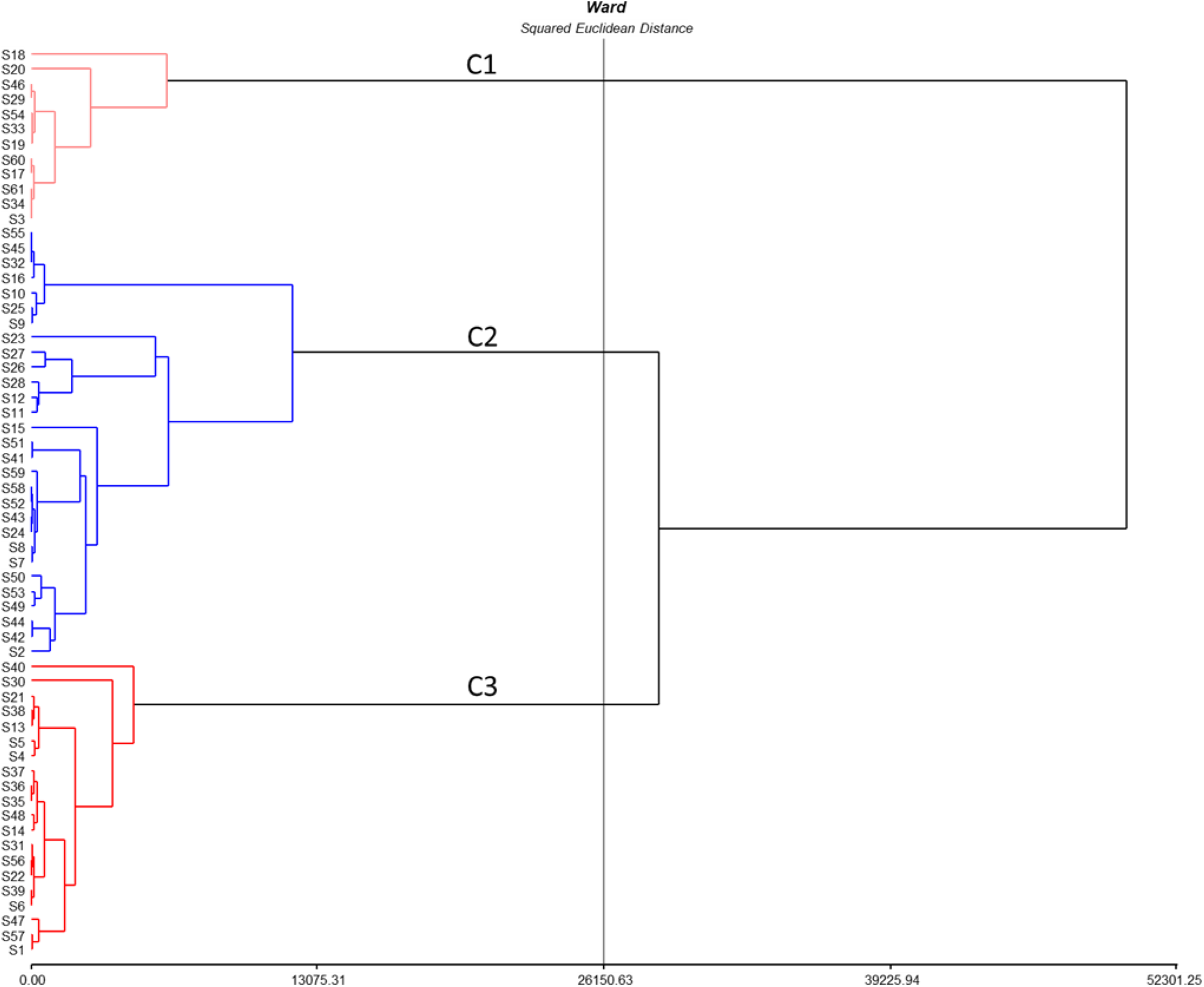
Cluster analysis of the microspheres with composition data by weight percentage (wt%) and diameters. Table 1. Chemical elements found in the microspheres (Semi-quantitative chemical analysis performed using a scanning electron microscope (SEM).

**Table 1.** Chemical elements found in the microspheres. (The chemical elements were obtained by semi-quantitative chemical analysis using a scanning electron microscope (SEM).

| Classification | Chemical elements |
| --- | --- |
| Alkali Metals | Na, K, Cs |
| Alkaline earth metals | Mg, Ca, Ba |
| Post transition metals | Al, Ga, In, Sn, Pb |
| Rare earth elements | La, Ce, Pr, Nd, Sm, Eu, Gd, Tb, Ho |
| **Transition metals** | **Ti, V, Mn, Fe, Co, Ni, Cu, Zn, Nb, Tc, Ru, Rh, Pd, Ag, Ta, Pt, Au** |
| **Metalloids** | **Si, As, Sb** |
| **Halogens** | **F, Cl, Br** |
| **Non metallic** | **C, N, O, P, S** |

The elements present in all microspheres were C and O (100%), followed by Si (92%), Al (78%), Fe (67%), and Ca (59%). Other elements were Mg, N, Ce, Ni, and La, identified in at least 36% of them. Less frequently, F and Na were present in at least 20% of the microspheres. The remaining elements were found in less than 10% of the microspheres, of which 14 elements were identified in a single sample (1.6%) (Table 2). For the highest weight percentage (%) of chemical elements detected present in at least one microsphere, there were Fe (92%), C (67%), O (52%), Ni (50%), Ce (48%), Pt (48%), Au (32%), La (31%), Si (27%), Ti (27%), Ca (20%), Ba (18%), F (12%), Al (11%) and N (11%) (Table 3). A characteristic feature of the microspheres was their size diversity from 1.3 µm to 32.5 µm (Table 4), indicating possible different development stages, and they were always isolated from themselves, although potential microorganisms were observed accompanying them.

**Table 2.** Percentages of total sampled microspheres containing a chemical element (CE). Semiquantitative chemical analysis performed using a scanning electron microscope (SEM) – EDS.

| CE | % | CE | % | CE | % | CE | % |
| --- | --- | --- | --- | --- | --- | --- | --- |
| C | 100.0 | F | 19.7 | Ag | 4.9 | Au | 1.6 |
| O | 100.0 | Ga | 9.8 | Pr | 4.9 | Cs | 1.6 |
| Si | 91.8 | P | 8.2 | S | 4.9 | As | 1.6 |
| Al | 78.7 | Co | 8.2 | Cl | 4.9 | Ta | 1.6 |
| Fe | 67.2 | Nd | 6.6 | Pb | 3.3 | Sm | 1.6 |
| Ca | 59.0 | V | 6.6 | Eu | 3.3 | Nb | 1.6 |
| Mg | 45.9 | Ba | 6.6 | Pd | 3.3 | Tc | 1.6 |
| N | 42.6 | In | 4.9 | Gd | 3.3 | Rh | 1.6 |
| Ce | 42.6 | Sb | 4.9 | Ru | 3.3 | Pt | 1.6 |
| Ni | 36.1 | Ti | 4.9 | Ho | 3.3 | Sn | 1.6 |
| La | 36.1 | Mn | 4.9 | Zn | 1.6 | K | 1.6 |
| Na | 19.7 | Tb | 4.9 | Cu | 1.6 | Br | 1.6 |

**Table 3.** Maximum percentages of chemical elements found in the estimated composition of the sampled microspheres. Semiquantitative chemical analysis performed using a scanning electron microscope (SEM) – EDS.

| CE | wt% | CE | wt% | CE | wt% | CE | wt% |
| --- | --- | --- | --- | --- | --- | --- | --- |
| Fe | 92.2 | F | 11.7 | Nd | 1.8 | Pb | 0.8 |
| C | 67.3 | Al | 11.4 | Ho | 1.8 | K | 0.8 |
| O | 52.3 | N | 10.7 | Tb | 1.7 | Cl | 0.7 |
| Ni | 49.8 | Ag | 5.9 | Ga | 1.6 | S | 0.7 |
| Ce | 48.3 | Ta | 5.6 | Pd | 1.4 | Rh | 0.6 |
| Pt | 48.2 | Cu | 4.2 | V | 1.3 | Sm | 0.6 |
| Au | 32.4 | Br | 2.8 | Eu | 1.2 | Cs | 0.5 |
| La | 31.1 | Mg | 2.8 | Sb | 1.1 | As | 0.4 |
| Si | 26.6 | In | 2.5 | Mn | 1.1 | Sn | 0.3 |
| Ti | 26.5 | Zn | 2.2 | Co | 1.1 | Ru | 0.3 |
| Ca | 19.5 | Gd | 2.0 | Pr | 1.0 | Nb | 0.3 |
| Ba | 17.7 | P | 1.8 | Na | 0.8 | Tc | 0.2 |

**Table 4.** General statistics and composition of the microspheres (% by weight – Wt%) in the most frequent chemical elements according to cluster (the analysis was based on the chemical elements determined in the microspheres and the chemical elements were obtained by semi-quantitative chemical analysis with a scanning electron microscope-SEM).

| cluster | statistic | $\varnothing (\mu\text{m})$ | C | O | Si | Al | Fe | Ca | Mg | N | Ce | Ni | La | Na | F | Ga | P | Co | Nd | V |
| --- | --- | --- | --- | --- | --- | --- | --- | --- | --- | --- | --- | --- | --- | --- | --- | --- | --- | --- | --- | --- |
| 1 | n | 12 | 12 | 12 | 12 | 12 | 12 | 12 | 12 | 12 | 12 | 12 | 12 | 12 | 12 | 12 | 12 | 12 | 12 | 12 |
|  | Media | 10.8 | 11.1 | 33.5 | 1.5 | 0.4 | 50.0 | 0.2 | 0.2 | 0.5 | 0.0 | 0.0 | 0.0 | 0.0 | 0.0 | 0.0 | 0.0 | 0.2 | 0.0 | 0.0 |
|  | D.E. | 5.2 | 3.0 | 13.6 | 1.3 | 0.6 | 14.8 | 0.5 | 0.7 | 1.1 | 0.0 | 0.0 | 0.0 | 0.0 | 0.0 | 0.0 | 0.0 | 0.4 | 0.0 | 0.1 |
|  | Min | 4.5 | 6.2 | 1.6 | 0.0 | 0.0 | 36.8 | 0.0 | 0.0 | 0.0 | 0.0 | 0.0 | 0.0 | 0.0 | 0.0 | 0.0 | 0.0 | 0.0 | 0.0 | 0.0 |
|  | Máx | 22.6 | 14.2 | 46.4 | 4.1 | 1.5 | 92.2 | 1.4 | 2.4 | 2.9 | 0.0 | 0.0 | 0.0 | 0.0 | 0.0 | 0.0 | 0.0 | 1.1 | 0.0 | 0.4 |
| 2 | n | 29 | 29 | 29 | 29 | 29 | 29 | 29 | 29 | 29 | 29 | 29 | 29 | 29 | 29 | 29 | 29 | 29 | 29 | 29 |
|  | Media | 7.7 | 38.9 | 29.6 | 8.5 | 2.3 | 6.4 | 2.8 | 0.8 | 2.9 | 2.4 | 0.2 | 0.2 | 0.2 | 0.0 | 0.0 | 0.1 | 0.0 | 0.1 | 0.1 |
|  | D.E. | 6.9 | 16.1 | 9.0 | 7.1 | 2.2 | 9.6 | 4.6 | 0.8 | 3.2 | 6.4 | 0.7 | 0.7 | 0.3 | 0.0 | 0.1 | 0.3 | 0.1 | 0.3 | 0.3 |
|  | Min | 1.3 | 9.4 | 12.4 | 0.7 | 0.0 | 0.0 | 0.0 | 0.0 | 0.0 | 0.0 | 0.0 | 0.0 | 0.0 | 0.0 | 0.0 | 0.0 | 0.0 | 0.0 | 0.0 |
|  | Máx | 32.5 | 67.3 | 52.3 | 26.6 | 11.4 | 38.8 | 19.5 | 2.8 | 10.7 | 27.7 | 3.4 | 3.0 | 0.8 | 0.0 | 0.3 | 1.8 | 0.5 | 1.8 | 1.3 |
| 3 | n | 20 | 20 | 20 | 20 | 20 | 20 | 20 | 20 | 20 | 20 | 20 | 20 | 20 | 20 | 20 | 20 | 20 | 20 | 20 |
|  | Media | 4.3 | 17.1 | 24.5 | 2.3 | 0.9 | 3.3 | 0.4 | 0.3 | 1.3 | 21.1 | 10.9 | 13.1 | 0.0 | 3.6 | 0.3 | 0.0 | 0.0 | 0.1 | 0.0 |
|  | D.E. | 2.1 | 7.0 | 7.5 | 1.6 | 0.7 | 4.2 | 0.4 | 0.5 | 1.9 | 9.5 | 9.8 | 6.6 | 0.1 | 3.4 | 0.6 | 0.1 | 0.0 | 0.3 | 0.0 |
|  | Min | 1.8 | 5.8 | 2.0 | 0.0 | 0.0 | 0.0 | 0.0 | 0.0 | 0.0 | 0.0 | 0.6 | 0.0 | 0.0 | 0.0 | 0.0 | 0.0 | 0.0 | 0.0 | 0.0 |
|  | Máx | 11.9 | 31.0 | 39.3 | 5.7 | 2.2 | 11.7 | 1.3 | 1.8 | 5.8 | 48.3 | 49.8 | 31.1 | 0.5 | 11.7 | 1.6 | 0.2 | 0.1 | 1.1 | 0.0 |
| All | n | 61 | 61 | 61 | 61 | 61 | 61 | 61 | 61 | 61 | 61 | 61 | 61 | 61 | 61 | 61 | 61 | 61 | 61 | 61 |
|  | mean | 7.2 | 26.3 | 28.7 | 5.1 | 1.5 | 14.0 | 1.5 | 0.5 | 1.9 | 8.1 | 3.7 | 4.4 | 0.1 | 1.2 | 0.1 | 0.1 | 0.0 | 0.1 | 0.0 |
|  | S.D. | 5.9 | 17.0 | 10.0 | 6.0 | 1.8 | 20.4 | 3.4 | 0.7 | 2.7 | 11.6 | 7.5 | 7.2 | 0.2 | 2.6 | 0.4 | 0.2 | 0.2 | 0.3 | 0.2 |
|  | min. | 1.3 | 5.8 | 1.6 | 0.0 | 0.0 | 0.0 | 0.0 | 0.0 | 0.0 | 0.0 | 0.0 | 0.0 | 0.0 | 0.0 | 0.0 | 0.0 | 0.0 | 0.0 | 0.0 |
|  | max. | 32.5 | 67.3 | 52.3 | 26.6 | 11.4 | 92.2 | 19.5 | 2.8 | 10.7 | 48.3 | 49.8 | 31.1 | 0.8 | 11.7 | 1.6 | 1.8 | 1.1 | 1.8 | 1.3 |

### Abundant Iron in the First Cluster

Cluster analysis could lead to the formation of possible groups that facilitate understanding this great diversity of independent microchondrules. Group 1 (C1) included twelve microspheres (Fig. 3), with an average diameter of 10.8 µm and average contents (Table 4) of C (11.1%), O (33.5%), and Fe (50.0%) in all microspheres (Figure 4); Si (1.5%) in nine; and N in three of them. Another notable aspect was the presence of two probable diatoms (Fig. 4). In this group, one microsphere stood out because, in addition to the four elements mentioned above, the presence of Ti (26.5%), Fe (44.4%) (Fig. 5) and other elements such as Ho (1.8%), Mg (2.4%), Al (1.3%) and Ca (1.4%) was evident, as well as V, Mn and N with less than 1%. Regarding the presence of Ti and V, it can be stated that it has a great coincidence with what was reported by Wainwright (2015), although other similar microspheres are presented later in this work.

**Fig. 4:**
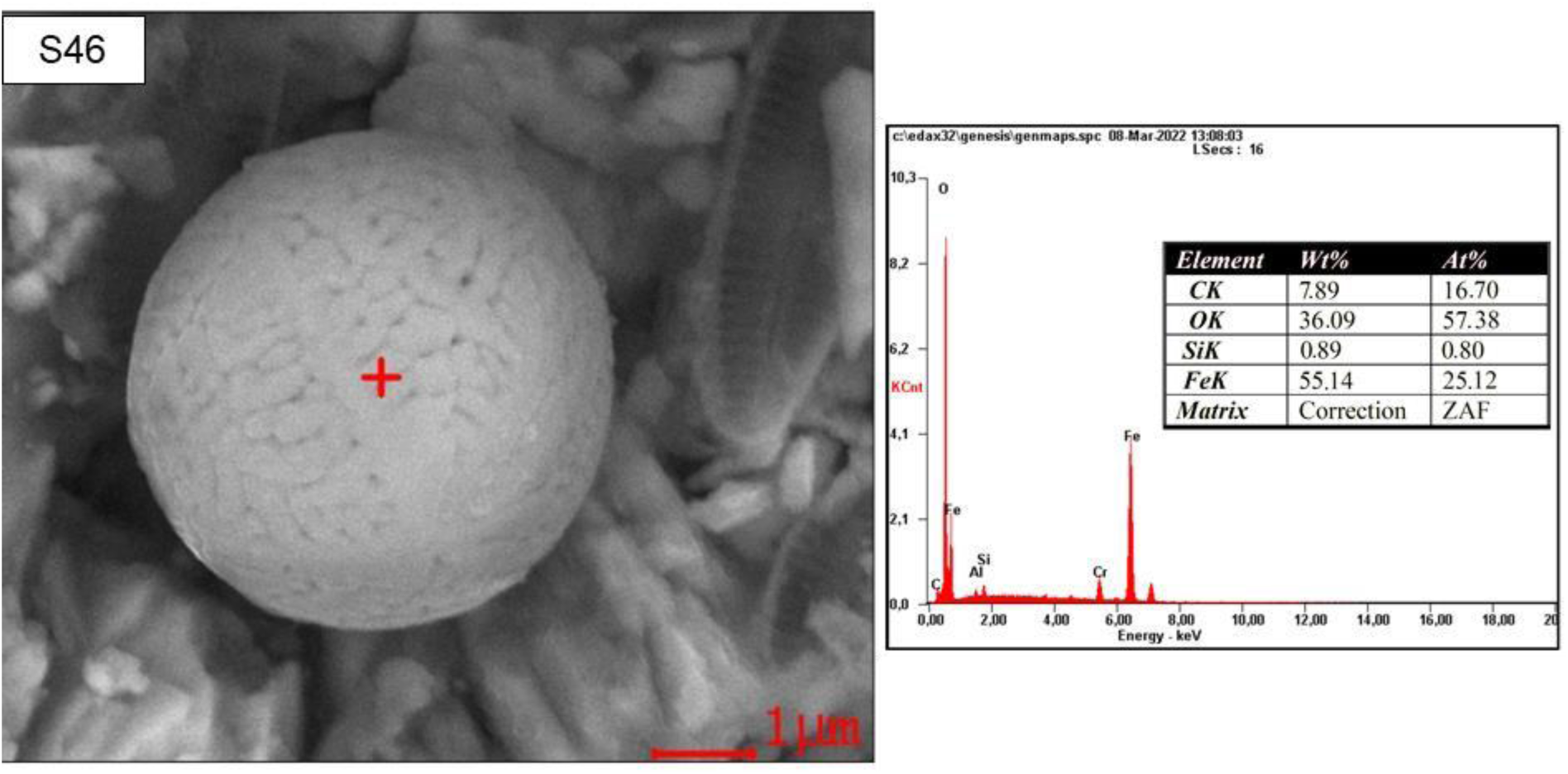
Microspheres containing primarily carbon, oxygen, and iron. Image obtained with a scanning electron microscope (SEM – EDS).

**Fig. 5:**
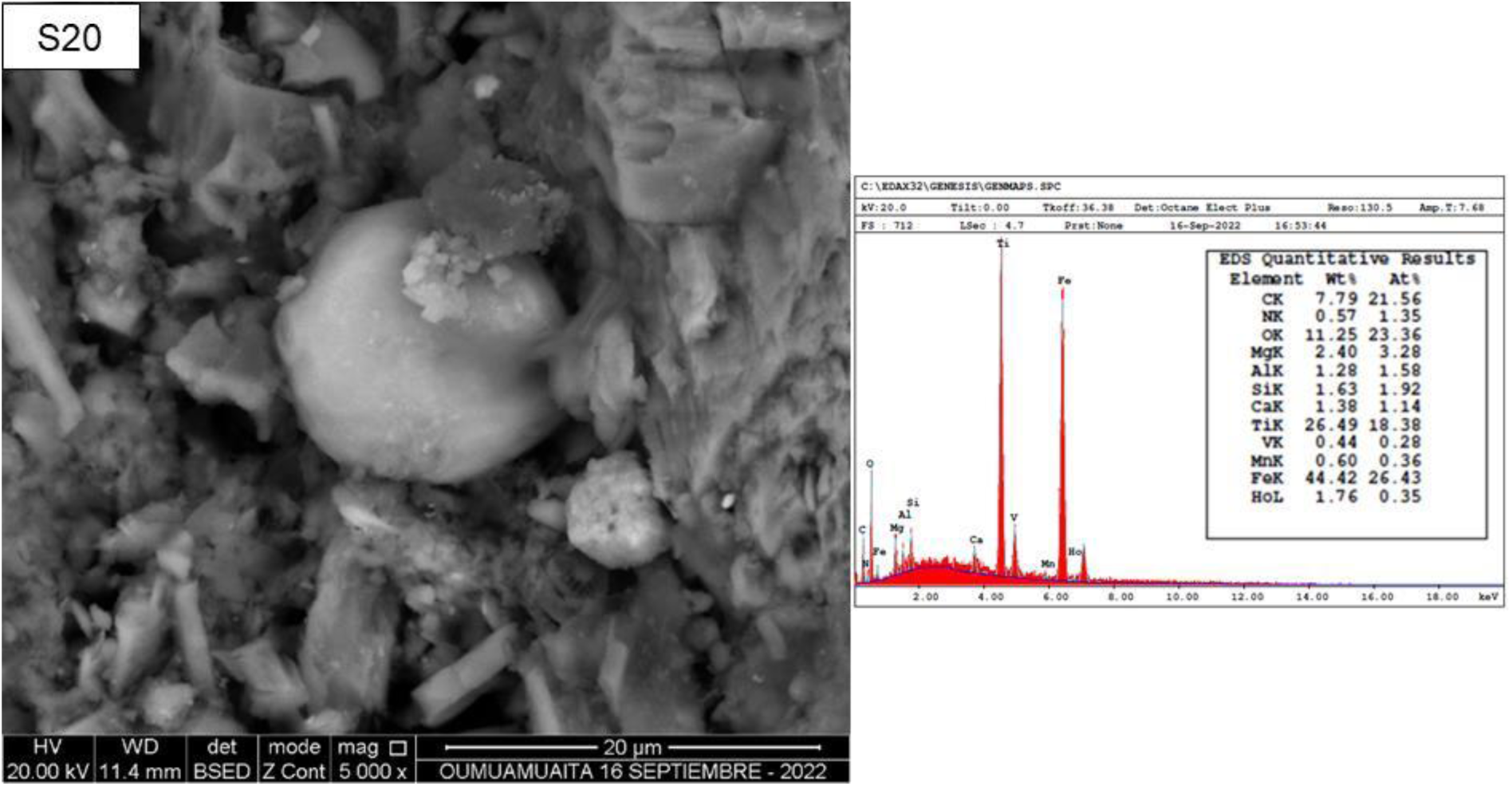
Microspheres containing primarily carbon, oxygen, iron, and titanium. Image obtained with a scanning electron microscope (SEM – EDS).

### Titanium, Vanadium, Silicon, and Aluminum in the Second Cluster

Cluster 2 (C2) contained twenty-nine microspheres (Fig. 2), with an average diameter of 7.7 µm and an estimated average content of C (38.9%), O (29.6%), and Si (8.5%). Other elements detected were Al (2.3%), Fe (6.4%), Ca (2.8%), Mg (0.8%), and N (2.9%) (Table 4) in at least 31% of the microspheres. In addition, a subgroup of seven microspheres was identified that, in addition to the aforementioned elements, contained Na, for a total of nine chemical elements.

In this cluster, two microspheres with Ti (between 1.5 and 15.1%) and V (between 0.2 and 3.9%) are notable (Fig. 6 and Fig. 7), similar to the 30 µm sphere described in the reviewed literature (Wainwright, 2015). Furthermore, one of them (Fig. 7) was similar in physical appearance and size, being 2.5 µm larger in diameter than the one observed in this study. A key difference worth highlighting is that the microspheres exist on the rock surface and were not the result of some type of microimpact, as was the case with the sphere recovered in Wainwright’s study (2015).

**Fig. 6:**
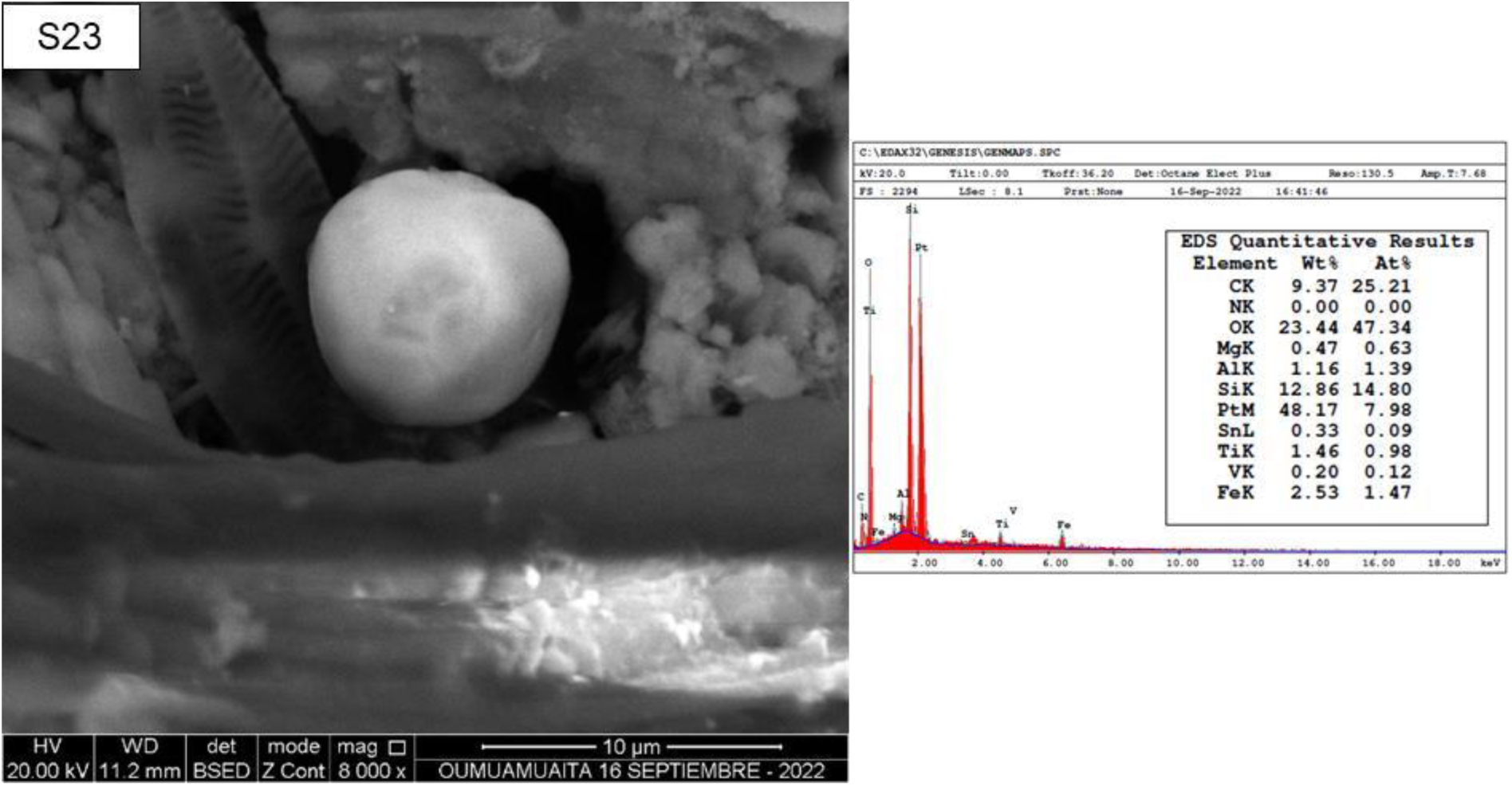
Microsphere containing primarily carbon, oxygen, silicon, and platinum, possibly accompanied by a diatom. Image obtained with a scanning electron microscope (SEM/EDS).

**Fig. 7:**
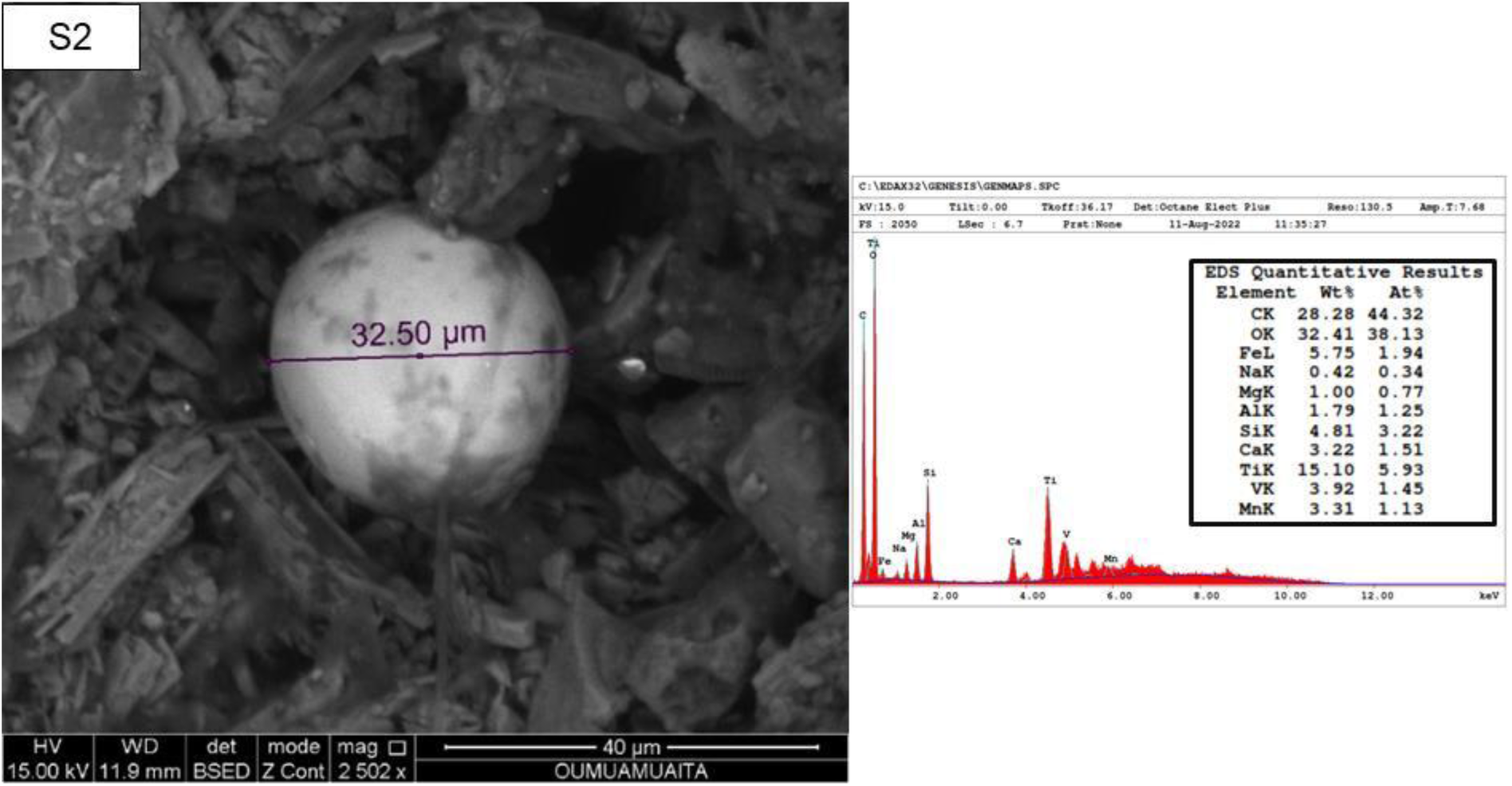
Microsphere composed primarily of carbon, oxygen, iron, titanium, and vanadium. Image obtained with a scanning electron microscope (SEM – EDS).

On the other hand, one of the microspheres from Cluster 2 also contained Mn (1.1%), Ba (17.7%), Eu (1.2%), Tb (1.4%) and Zn (2.2%), while the other stood out for being the only one with contents of Pt (48.2%) and Sn (Fig. 6). A notable fact when observing the microsphere with high Pt content was the presence of an organism, apparently diatom-like, that accompanied the sphere and was in very good condition, giving the appearance of being alive; similar to what is shown in Fig. 4, although it is well known that it is not possible to observe living organisms in a SEM.

Furthermore, this cluster contained the only sphere with Au content (between 27.2 and 37.6%) that was identified in the presence of other elements, such as Ag (between 4.1% and 7.7%), Cu (between 3.7% and 4.8%) (Fig. 8), Pr, Cs, Tb and Mn. Finally, the presence of other non-metallic elements, such as P, Cl, Br and S; metallic elements (Ni, Pd, K, Rh, In and Co); and rare earth elements (La, Ce and Nd) were detected in some microspheres.

**Fig. 8:**
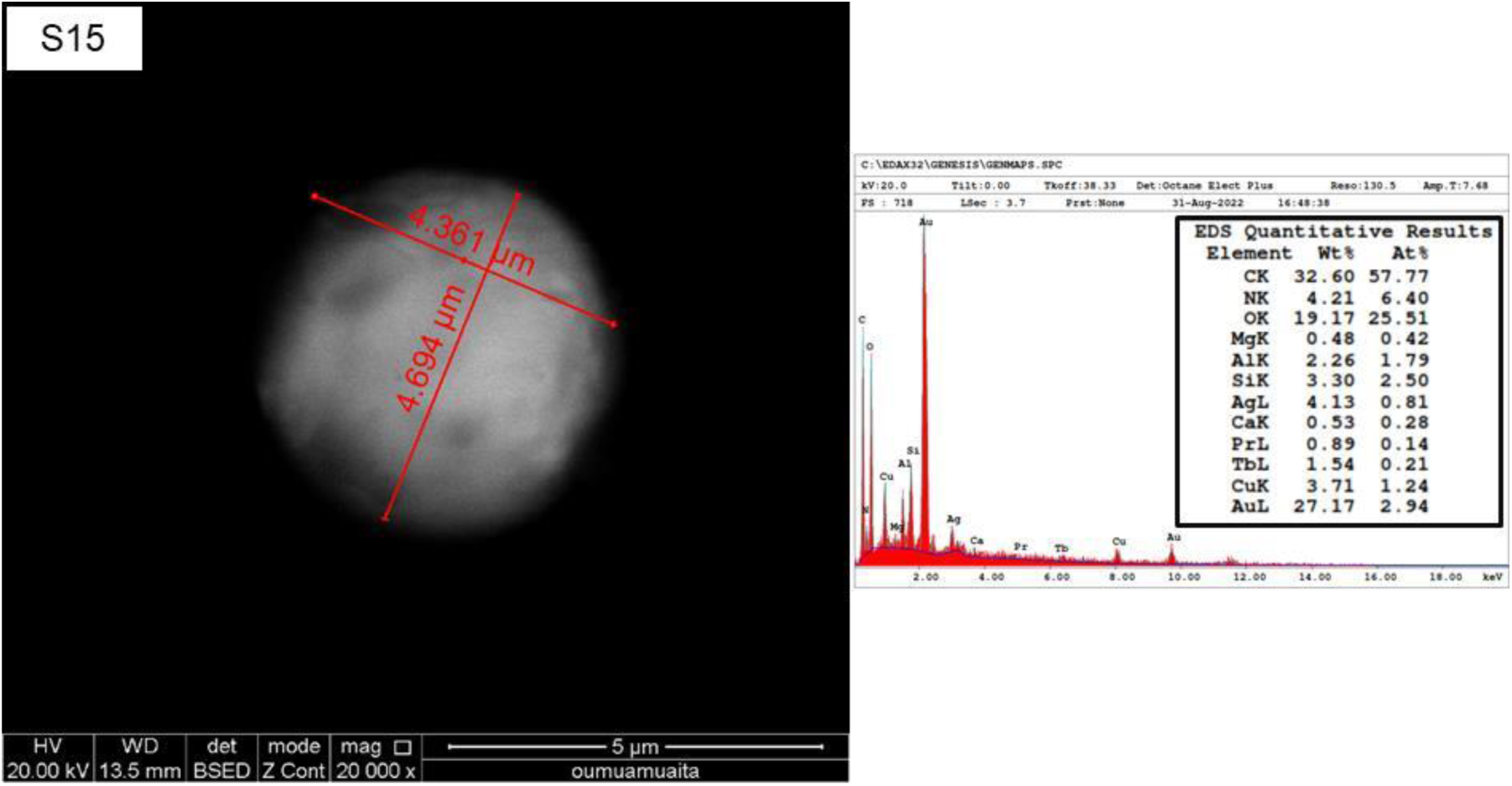
Microspheres containing primarily carbon, oxygen, and gold. Image obtained with a scanning electron microscope (SEM – EDS).

### Cerium, Lanthanum, and Nickel in the Third Group

Group 3 consisted of twenty microspheres (Fig. 2), with an average diameter of 4.3 µm. The main characteristics were the presence of C (17.1%), O (24.5%), and Ni (10.9%) in all of them, as well as La (13.1%) and Ce (21.1%) in nineteen of them, Si (2.3%) in eighteen, Al in sixteen, F (3.6%) in twelve, Ca in eleven, Fe (3.3%) in ten, N (1.4%) and Mg in seven of them (Table 4). In addition to these elements, with the exception of F, one microsphere was found with the presence of Na, P, Pb, and Co, for a total of fifteen elements. Some microspheres in the group contained metallic elements, In, Pb, Co, Nd, and Ag, and one metalloid, Sb. In addition, there was one sphere containing As, Ta, Pd, Sm, and Gd; another sphere containing S, Tc, and Ru; and one microsphere containing twelve chemical elements, including Nd (1.1%) and Nb (Fig. 9).

**Fig. 9:**
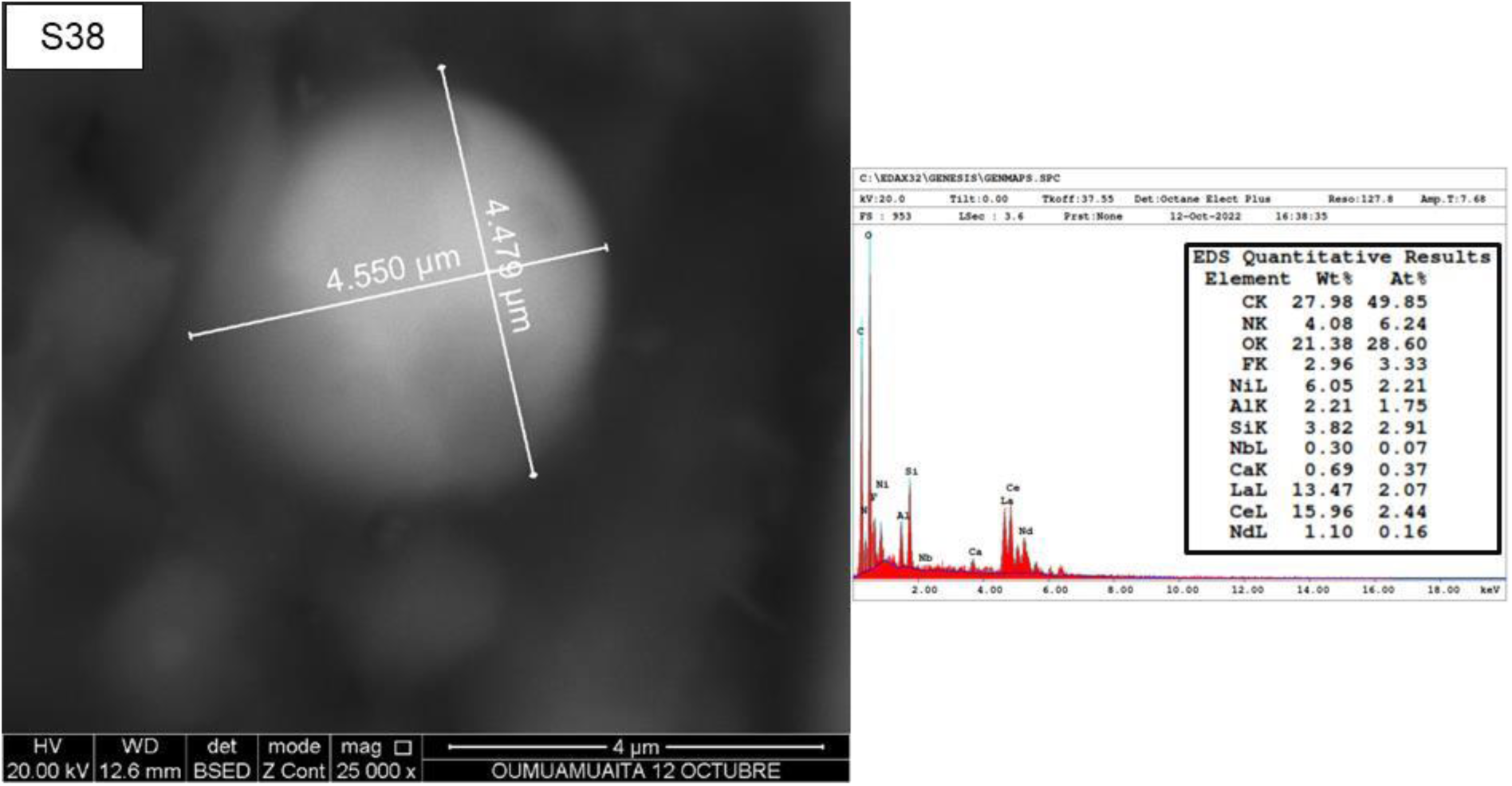
Microspheres containing primarily carbon, oxygen, lanthanum, cerium, and nickel. Image obtained with a scanning electron microscope (SEM – EDS).

### Analysis of microspheres according to morphological characteristics

From the images corresponding to the large numbers of BEs, four clusters were identified based on their morphology (M1, M2, M3, and M4) that exhibit different characteristics (Fig. 10).

**Fig. 10:**
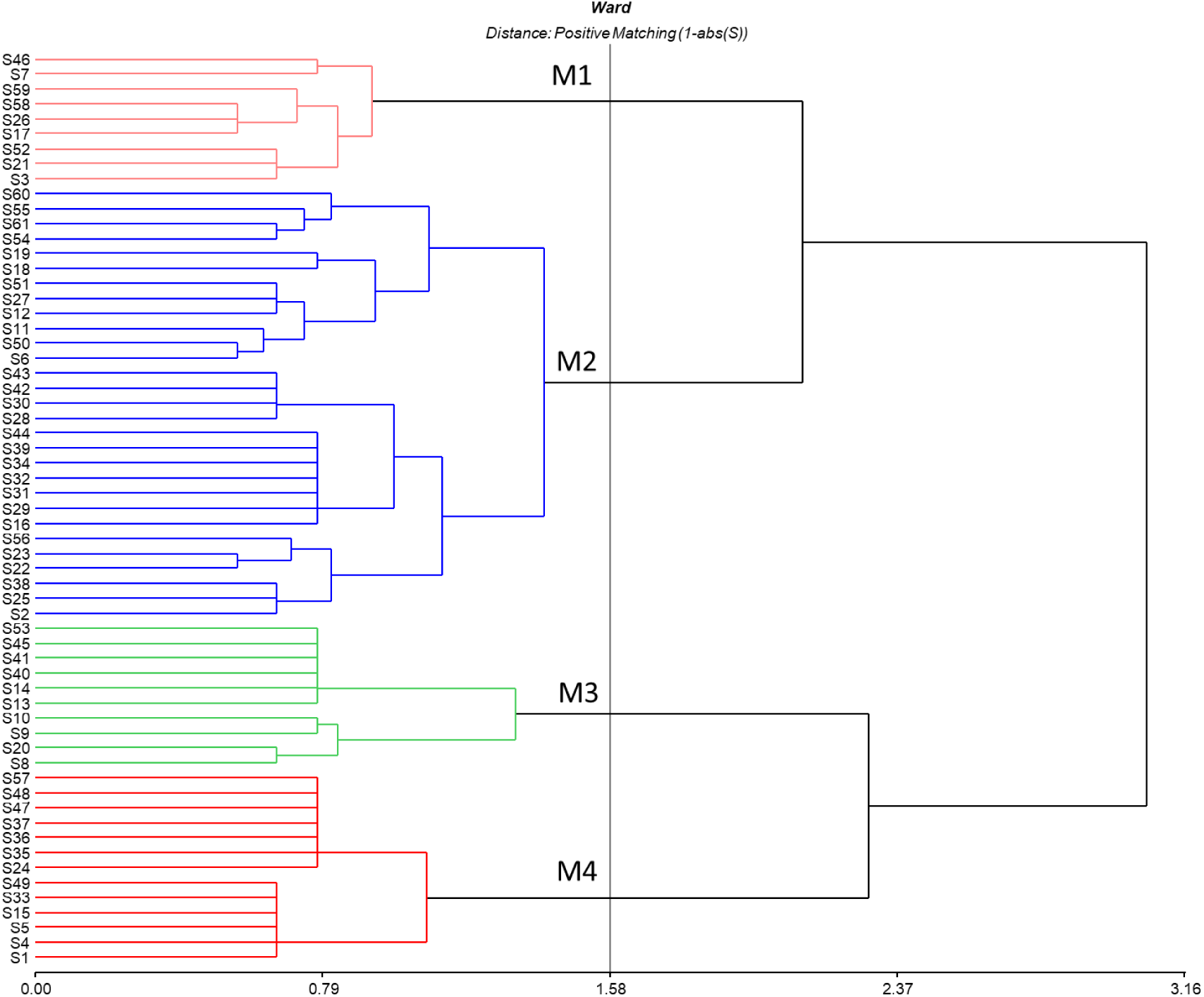
Cluster analysis of the microspheres with morphometric data (microsphere shapes). Semiquantitative chemical analysis performed using a scanning electron microscope (SEM) – EDS.

### Brain-like Microspheres

Group 1 (M1) contains nine brain-like microspheres; they are spherical, some with spots and scales, and range in size from small to large (Fig. 11). These microspheres were characterized by the presence of C, O, and Si; Fe was also found in six of them, and N, Al, Na, Mg, and Ca in one. These microspheres had a longitudinal structure that joined two hemispheres and was different from the fissure characteristic of a human brain. In this group, the microchondrules ranged in size, from 9.9 µm to 22.6 µm; a higher abundance of iron and a reduction in carbon content were observed, as well as the absence of some elements in the larger ones, which may be evidence of the existence of different developmental stages and a particular species of brain-like spherical microchondrules (Fig. 11).

**Fig. 11:**
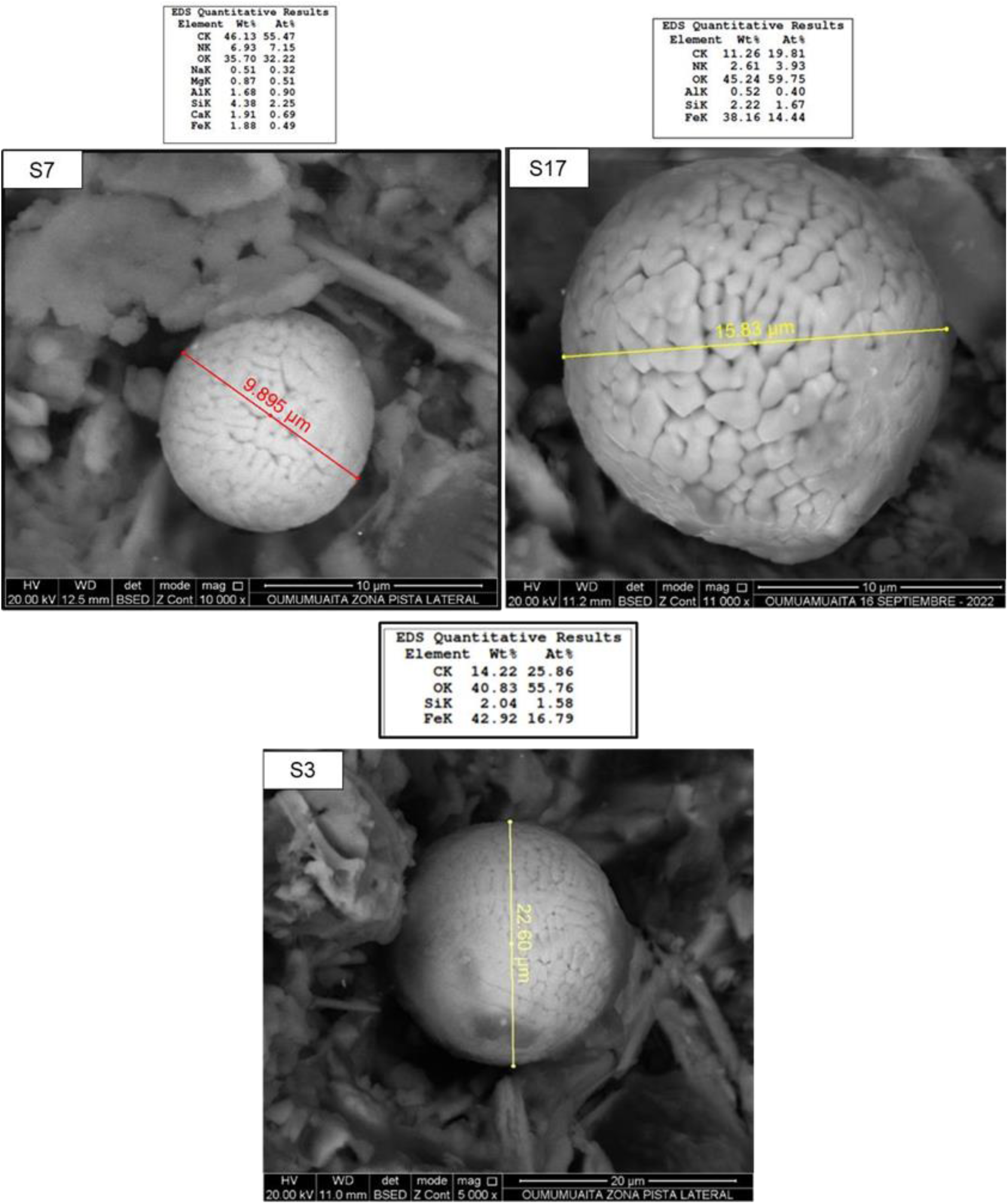
Brain-like microspheres of varying sizes and compositions (Rock in Colombia). Semiquantitative chemical analysis performed using a scanning electron microscope (SEM) – EDS.

One behavior observed in these microspheres is their displacement. Several attempts have been made to perform chemical analysis of brain-like microspheres using a scanning electron microscope (SEM), but in ten analysis attempts, the microspheres moved or disappeared, preventing scanning. Figure 12 shows this phenomenon in real time. The first image identifies the area where the microsphere is located (Figure 12a), then the image zooms in to identify the central object (Figure 12b), at which point the brain-like microsphere disappears from where it was located (Figures 12c and 12d), and the image of the general study area is returned (Figures 12e and 12f).

**Fig. 12.**
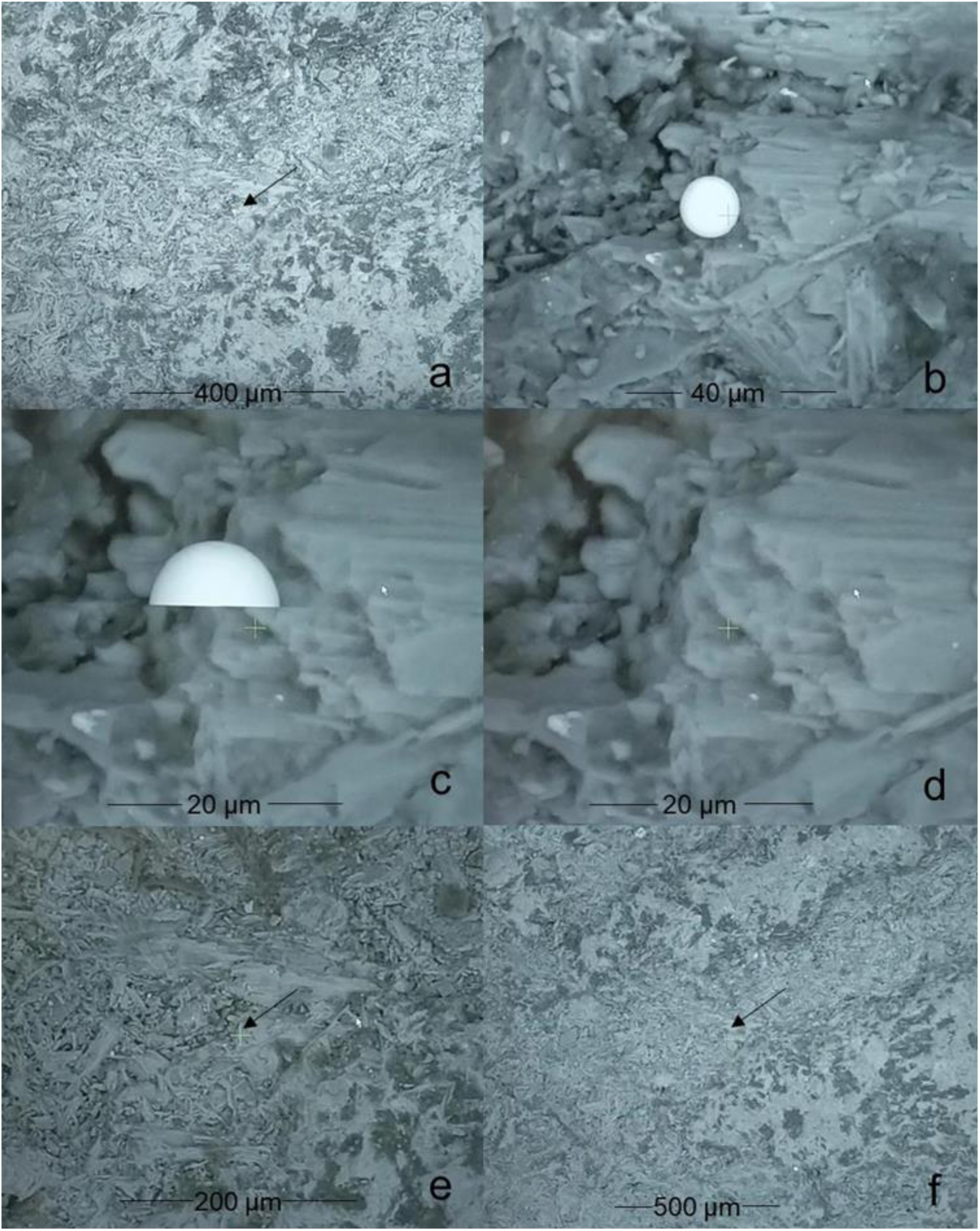
Movement of a brain-like microsphere during scanning for analysis. Semiquantitative chemical analysis performed using a scanning electron microscope (SEM-EDS).

**Fig. 12:**
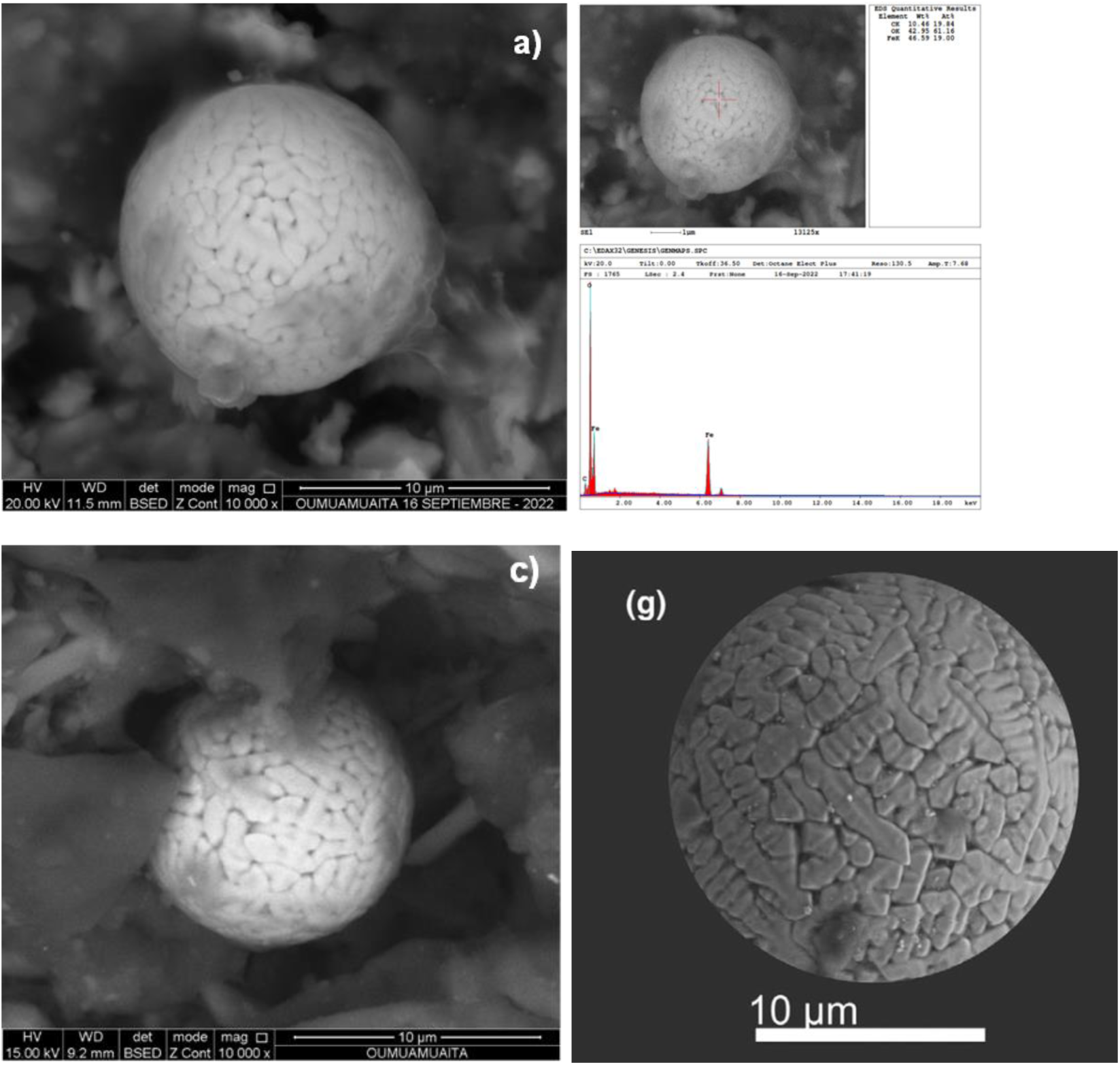
a) and c) Brain-like microspheres composed of iron oxide and carbon found in the rock and b) semi-quantitative chemical analysis. Semi-quantitative chemical analysis performed using a scanning electron microscope (SEM – EDS). (g) Dust particles obtained by filtering fresh snow collected from May to September 2017 in the vicinity of Vostok Station, Antarctica were examined using a scanning electron microscope. The dust particle collection contains 197 spherules ranging from 0.5 to 117 μm in diameter, with iron oxide spherules being by far the most abundant (n = 188). Analysis of Meteorological and human activity data suggest an extraterrestrial origin for most of the spherical particles.

**Figure 13:**
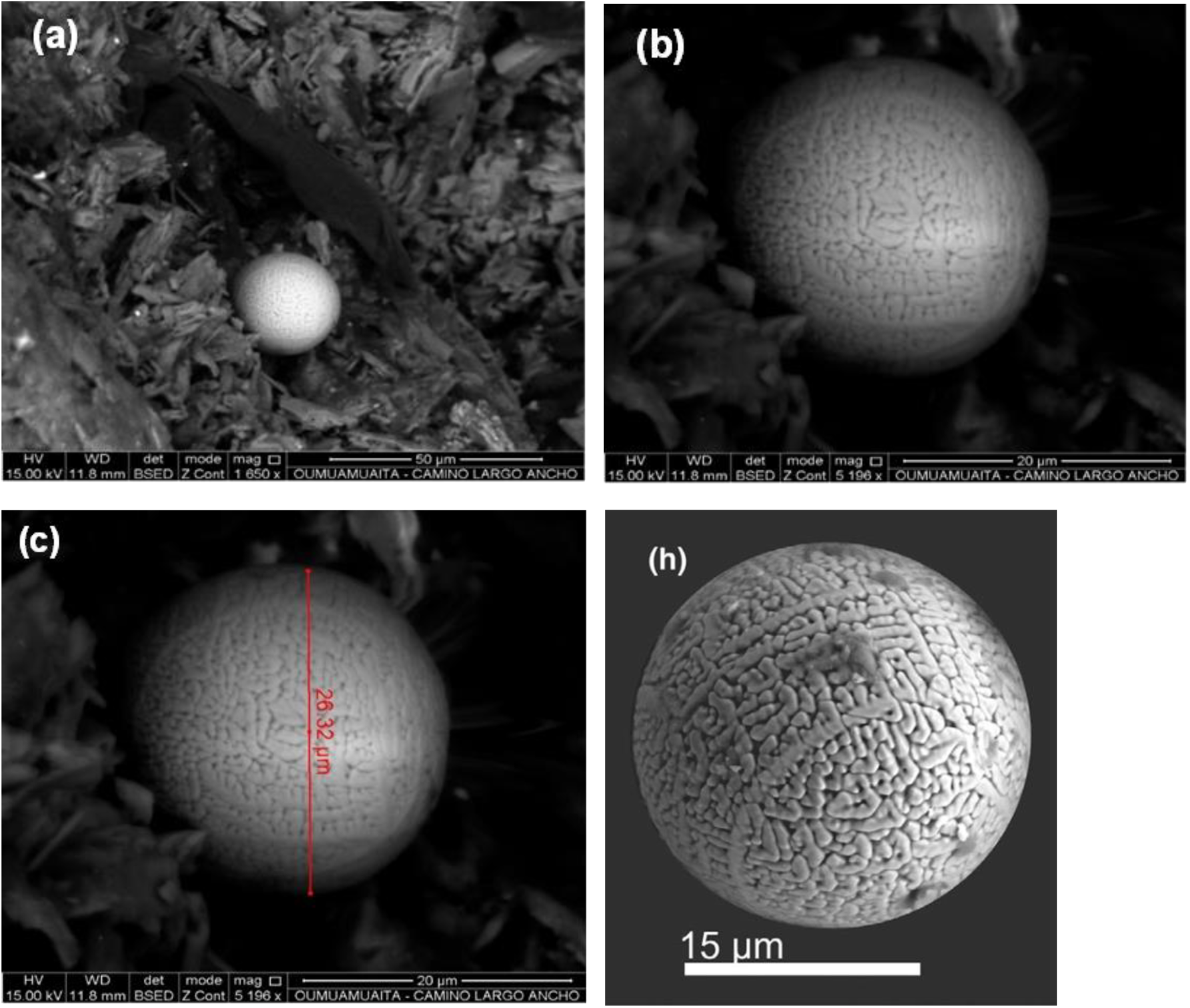
Microsphere found in LA ROCA – August 2023. Microimages (a), (b) and (c), respectively. Analysis performed on April 13, 2023. Semi-quantitative chemical analysis performed with a scanning electron microscope (SEM – EDS). Image (h). Dust particles obtained by filtering fresh snow collected from May to September 2017 near Vostok Station in Antarctica were examined using a scanning electron microscope. The dust particle collection contains 197 spherules ranging in diameter from 0.5 to 117 μm, with iron oxide spherules being by far the most abundant (n = 188). Analysis of meteorological and human activity data suggests an extraterrestrial origin for most of the spherical particles.

**Figure 14:**
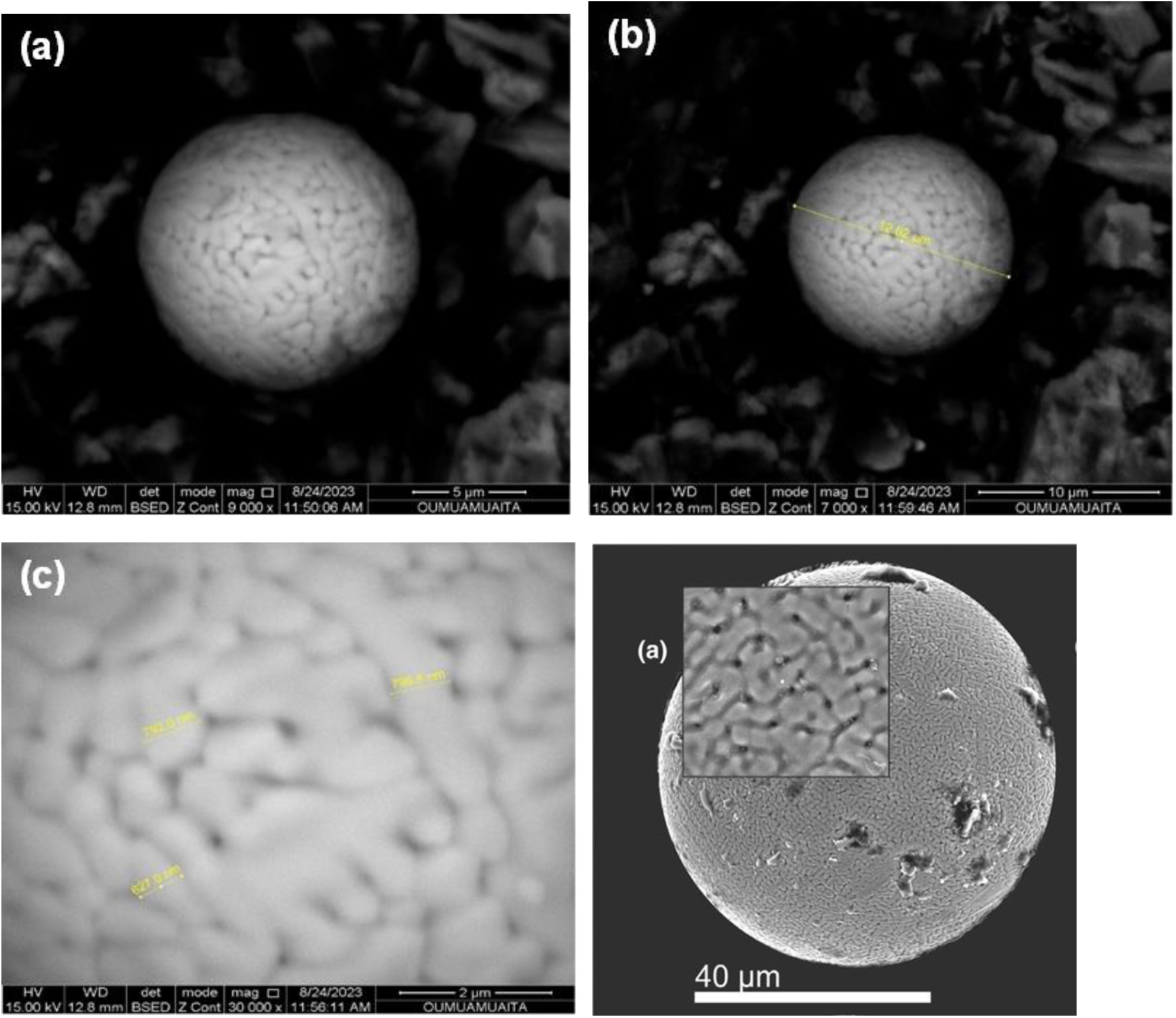
Microsphere found in LA ROCA – August 2023. Microimages (a), (b) and (c), respectively. Semi-quantitative chemical analysis performed with a scanning electron microscope (SEM – EDS). Image (1), (a). Dust particles obtained by filtering fresh snow collected from May to September 2017 in the vicinity of Vostok Station in Antarctica were examined using a scanning electron microscope. The dust particle collection contains 197 spherules ranging from 0.5 to 117 μm in diameter, with iron oxide spherules by far the most abundant (n = 188). Analysis of meteorological and human activity data suggests an extraterrestrial origin for most of the spherical particles. Source: https://new.ras.ru/activities/news/kosmicheskaya-pyl-v-sugrobakh-antarktidy/, 2023.

**Figure 15.**
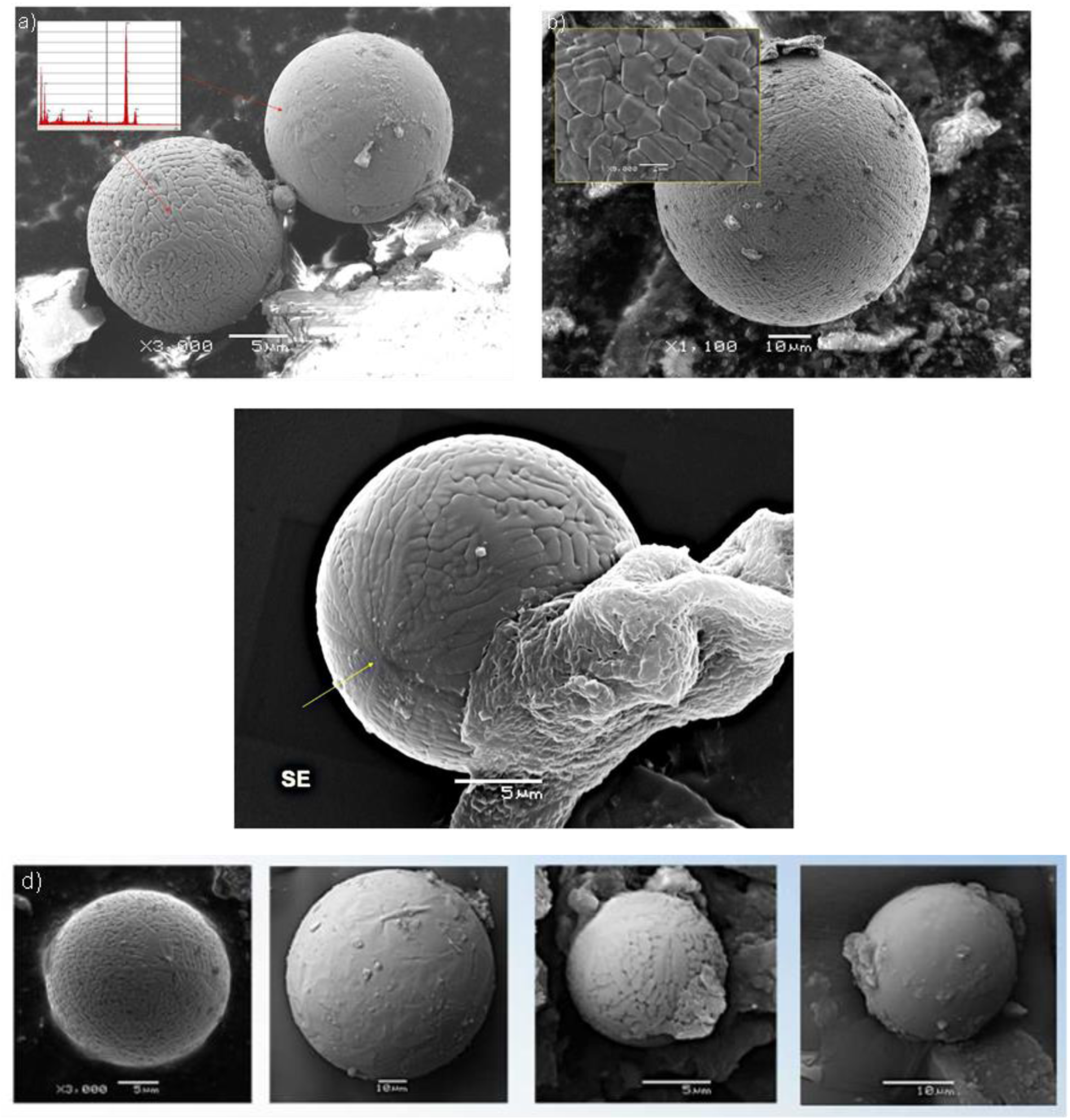
Microspherules found near the Kamil crater: Southwest of New Valley Governorate, Egypt. Note the similarity in morphology and chemical composition with the microspheres found in THE ROCK, Figures 4 and 12 (a) of this document. Source: https://www.researchgate.net/publication/281787526-2014.

**Figure 16.**
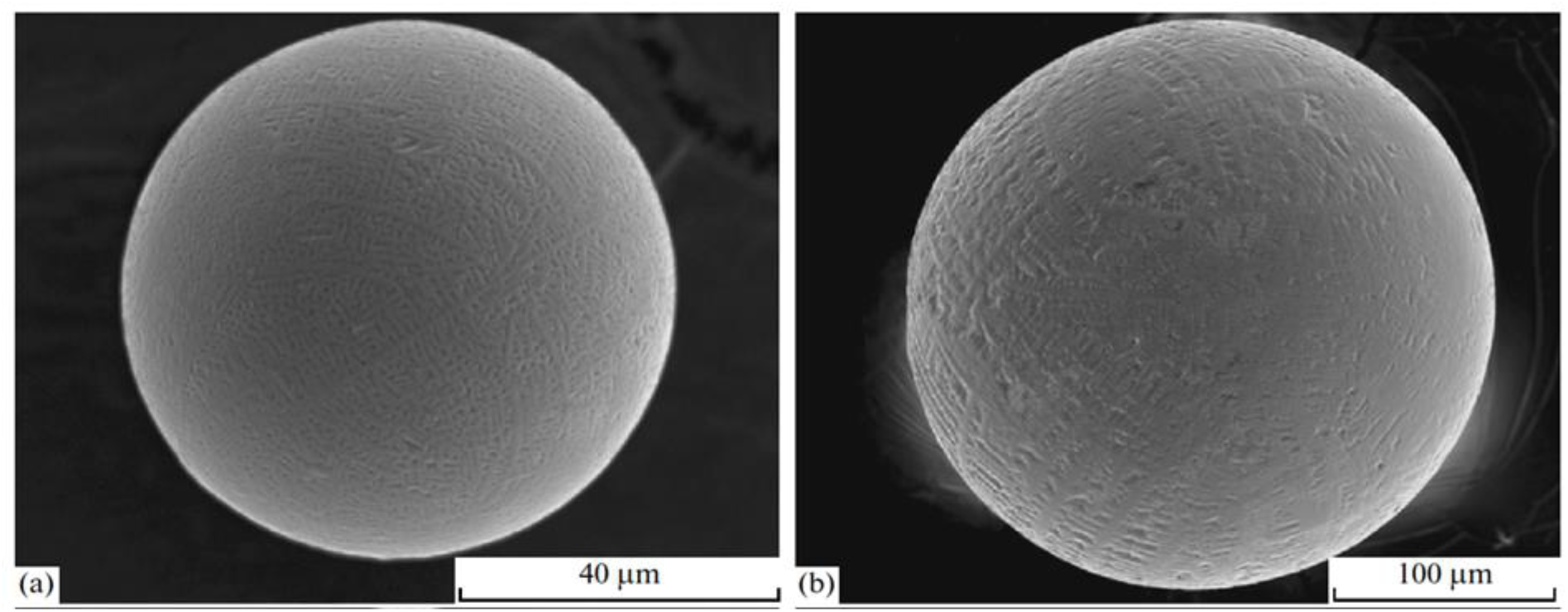
Microspherules found in the Sikhote Alin meteorite. Note the similarity in morphology and chemical composition with the microspheres found in LA ROCA, Figures 13 (c) of this document. Source: https://www.researchgate.net/publication/257847393 2012.

**Figure 17.**
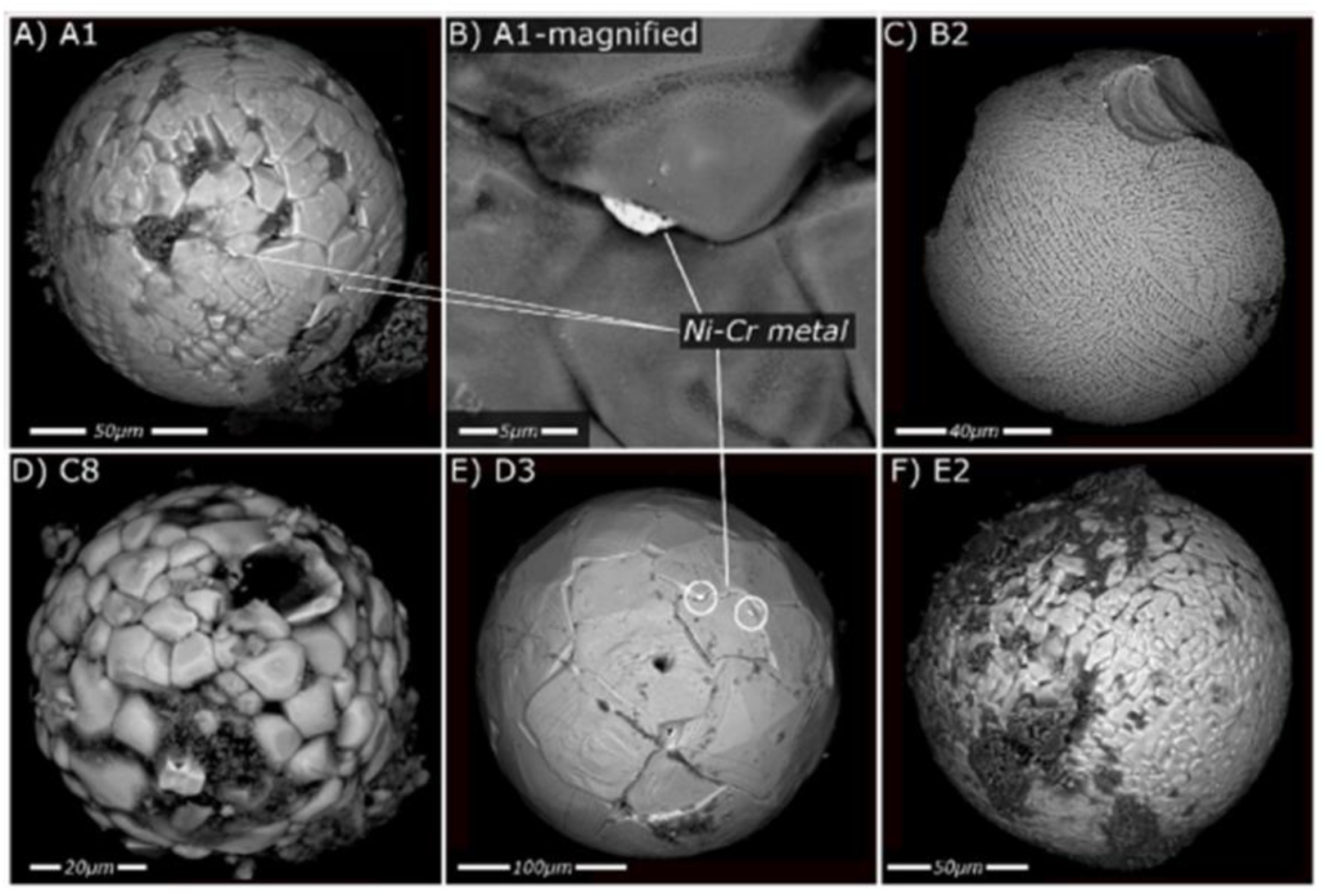
Fossil micrometeorites from Monte dei Corvi found in sediments, searching for dust from the Veritas asteroid family and the usefulness of micrometeorites as a paleoclimate proxy. Note the similarity (C) B2, in morphology and chemical composition, with the microspheres found in LA ROCA, Figure 13 (c), of this document. Source: https://www.sciencedirect.com/science/article/pii/S0016703723003010?via%3Dihub 2023.

**Figure 18.**
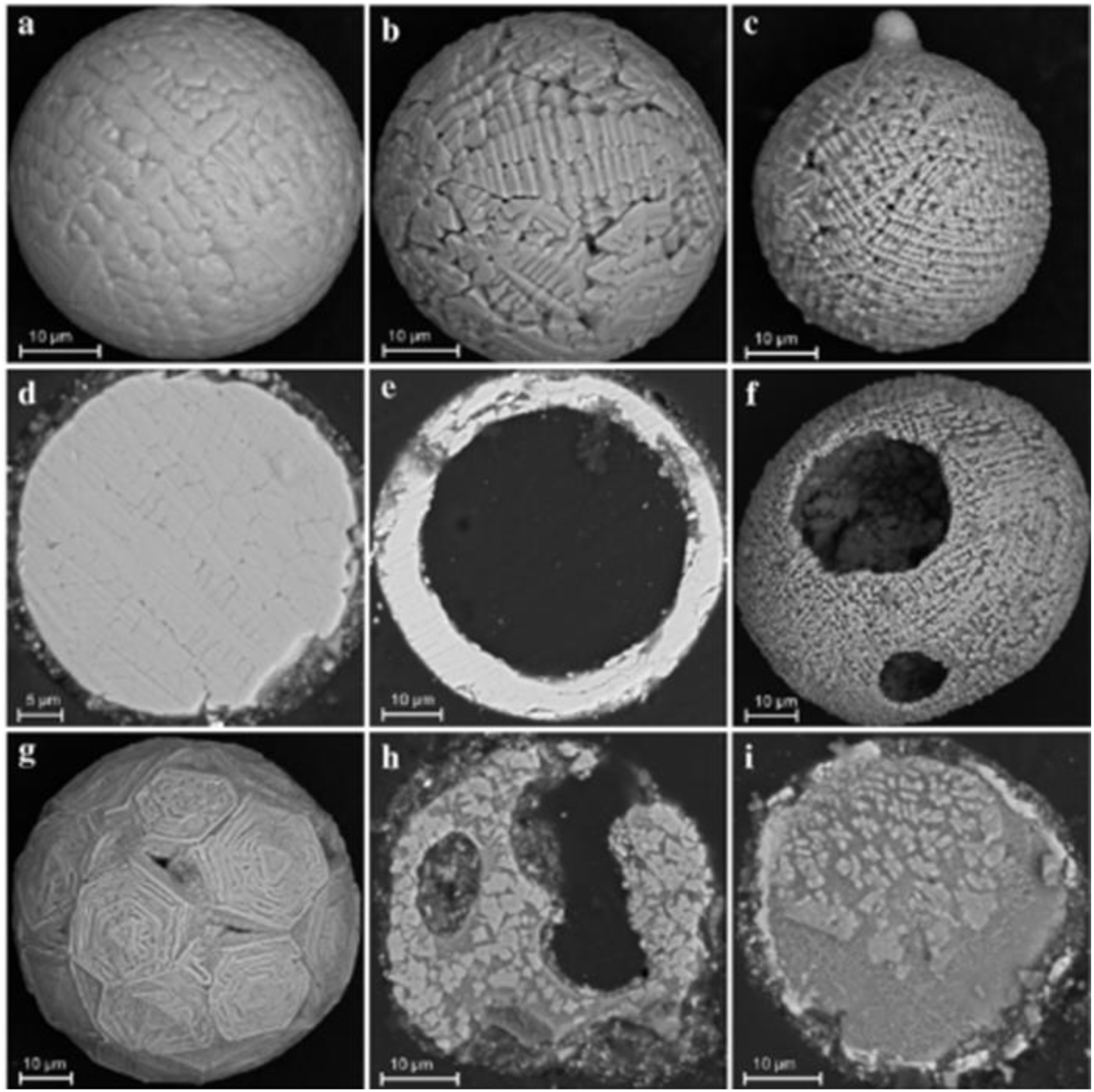
Microspheres from a Tunguska meteorite explosion in 1908, Siberia, Russia. Note the similarity in morphology and chemical composition to the microspheres found in LA ROCA (Rock), Figure 14 (a), in this document. Source: https://piramidesdebosnia.com/2017/08/20/misterios-del-valle-de-la-muerte-parte-4-ultima/ 2023

**Figure 19:**
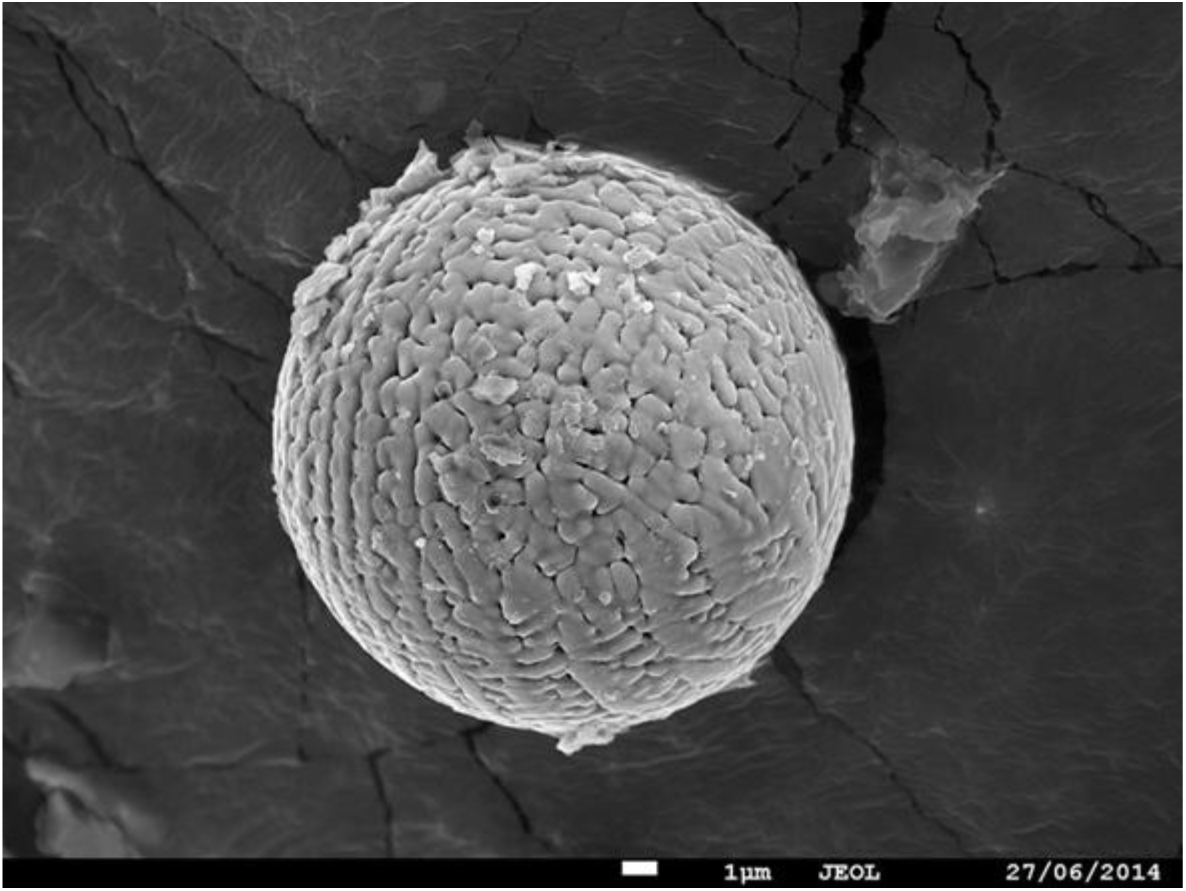
Minimeteorites A 2.7 billion-year-old micrometeorite extracted from limestone rock found in the Pilbara region of Western Australia. Monash University, Australia. Composed of iron oxide. Source: Proceedings of the National Academy of Sciences, January 21, 2020.

**Figure 20:**
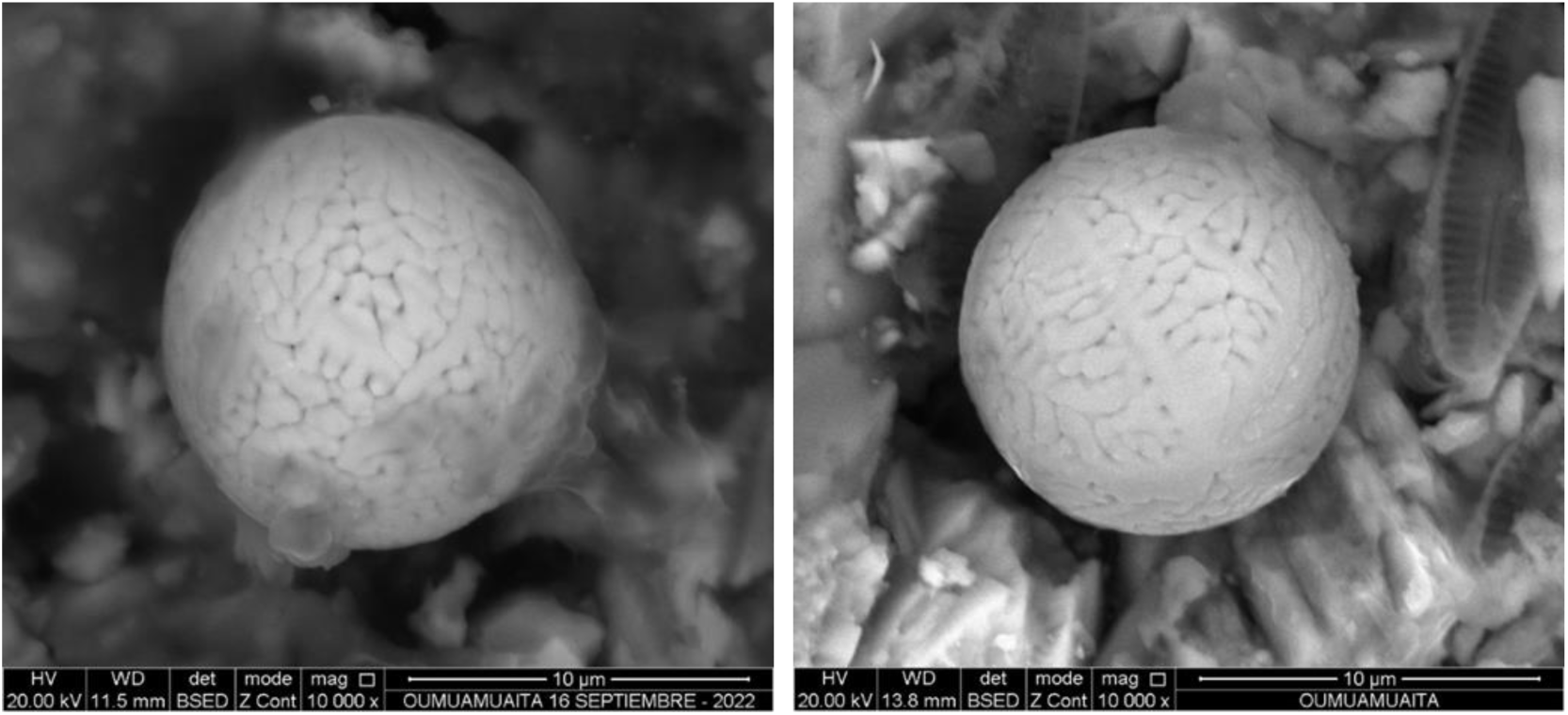
Microspheres found in LA ROCA – (a) analysis carried out on September 16, 2022 and (b) analysis carried out on March 8, 2022. Note the similarity and equality to the previous image. Semi-quantitative chemical analysis carried out with a scanning electron microscope SEM – EDS.

**Figure 21.**
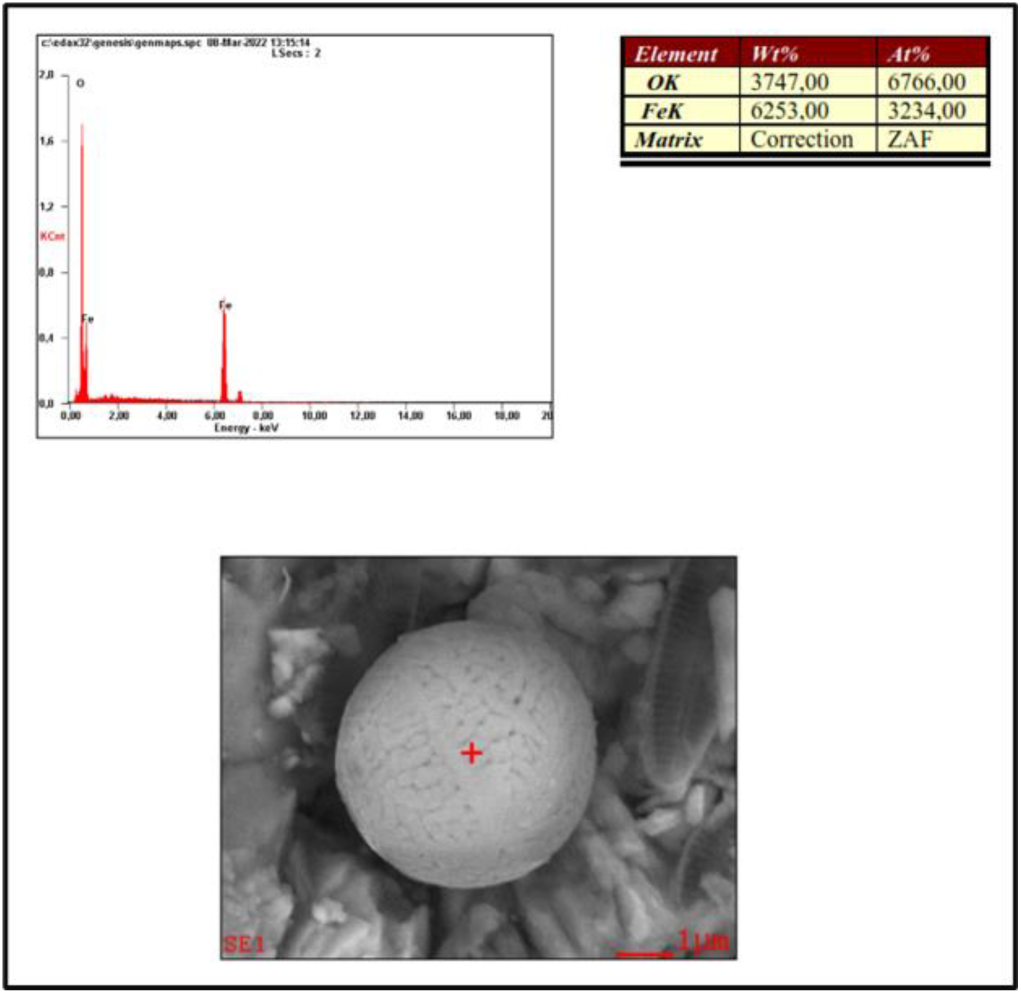
Elemental chemical analysis of the brain-like microsphere: Semi-quantitative chemical analysis performed using a scanning electron microscope (SEM) – EDS. Note the identical composition to the micrometeorite found in Australia: Figure 19.

**Figure 22.**
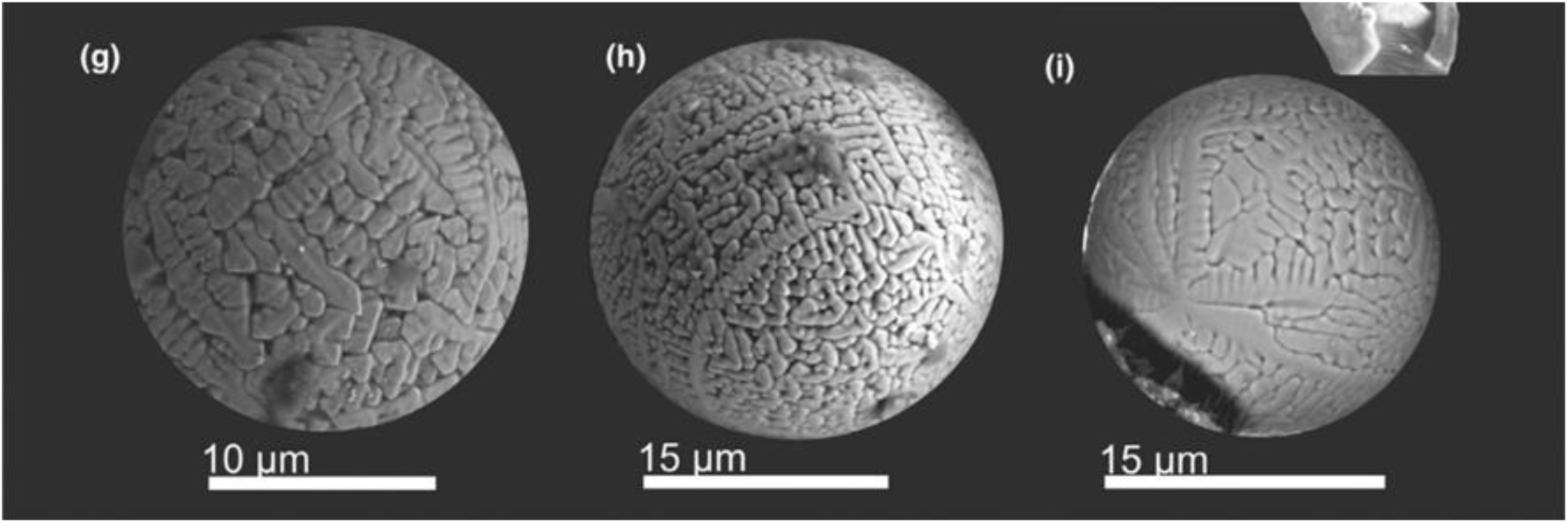
Images (g), (h), and (i). Dust particles obtained by filtering fresh snow collected from May to September 2017 in the vicinity of Vostok Station in Antarctica were examined using a scanning electron microscope. The dust particle collection contains 197 spherules ranging from 0.5 to 117 μm in diameter, with iron oxide spherules by far the most abundant (n = 188). Analysis of meteorological and human activity data suggests an extraterrestrial origin for most of the spherical particles. Source: https://new.ras.ru/activities/news/kosmicheskaya-pyl-v-sugrobakh-antarktidy/ 2023. Image (h)

**Figure 23.**
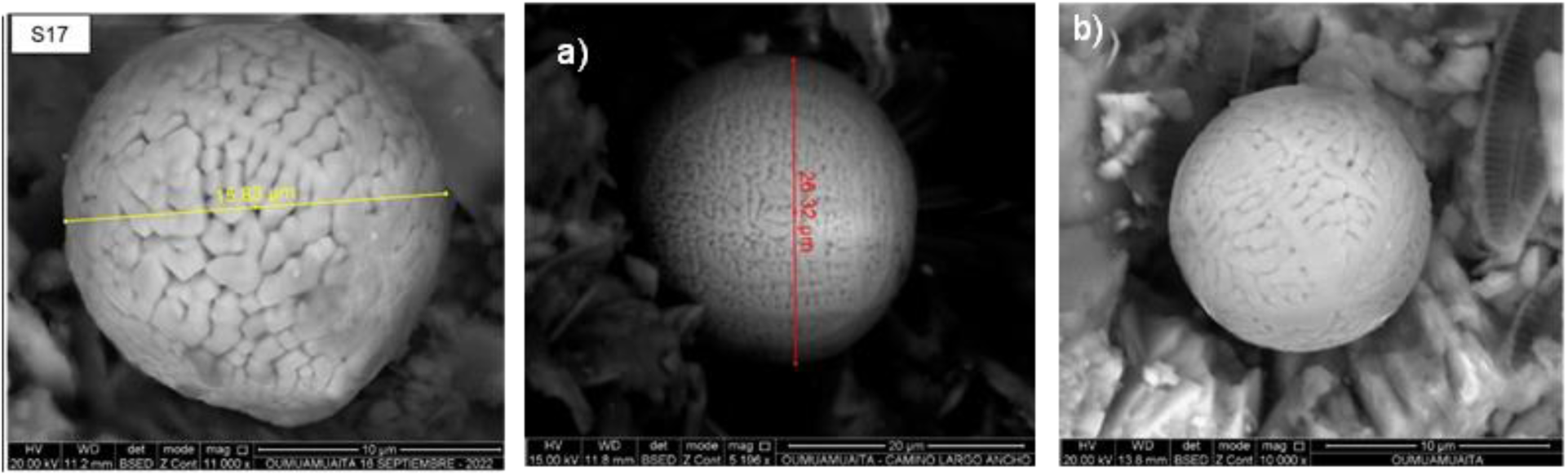
Images (S17), (a) and (b), microspheres found at LA ROCA (2002 and 2023, respectively); note the similarity with the microspherules collected from May to September 2017 in the vicinity of Vostok Station in Antarctica and examined using a scanning electron microscope. Source: Own, images obtained with a scanning electron microscope SEM – EDS.

In an analysis in scanning electron microscope SEM - EDS, performed on a microspherule located in the rock of the brain-type class gave a different elemental composition (Figure 24 a), to the average compositions of the brain-type spherules analyzed previously (Figure 24 b), no iron Fe was found, but other elements totally different from those found in the previous microspherules, Nickel (Ni), Aluminum (Al), Tantalum (Ta), Lanthanum (La), Cerium (Ce) and Neodymium (Nd), this microspherule was taken from the rock and placed on a graphene PIN, it was analyzed in a dual beam SEM - EDS scanning electron microscope, finding the same composition found in the same microspherule in the rock (Figure 25). Subsequently, another elemental chemical analysis was performed using a backscattered scanning electron microscope (SEM-EDS) at the Geological Institute of Mexico at the National Autonomous University of Mexico (UNAM). It was found that its composition had changed slightly. New elements such as vanadium (V), fluorine (F), barium (Ba), and cesium (Cs) were found, but no cerium (Ce) was found (Figure 26).

**Figure 24.**
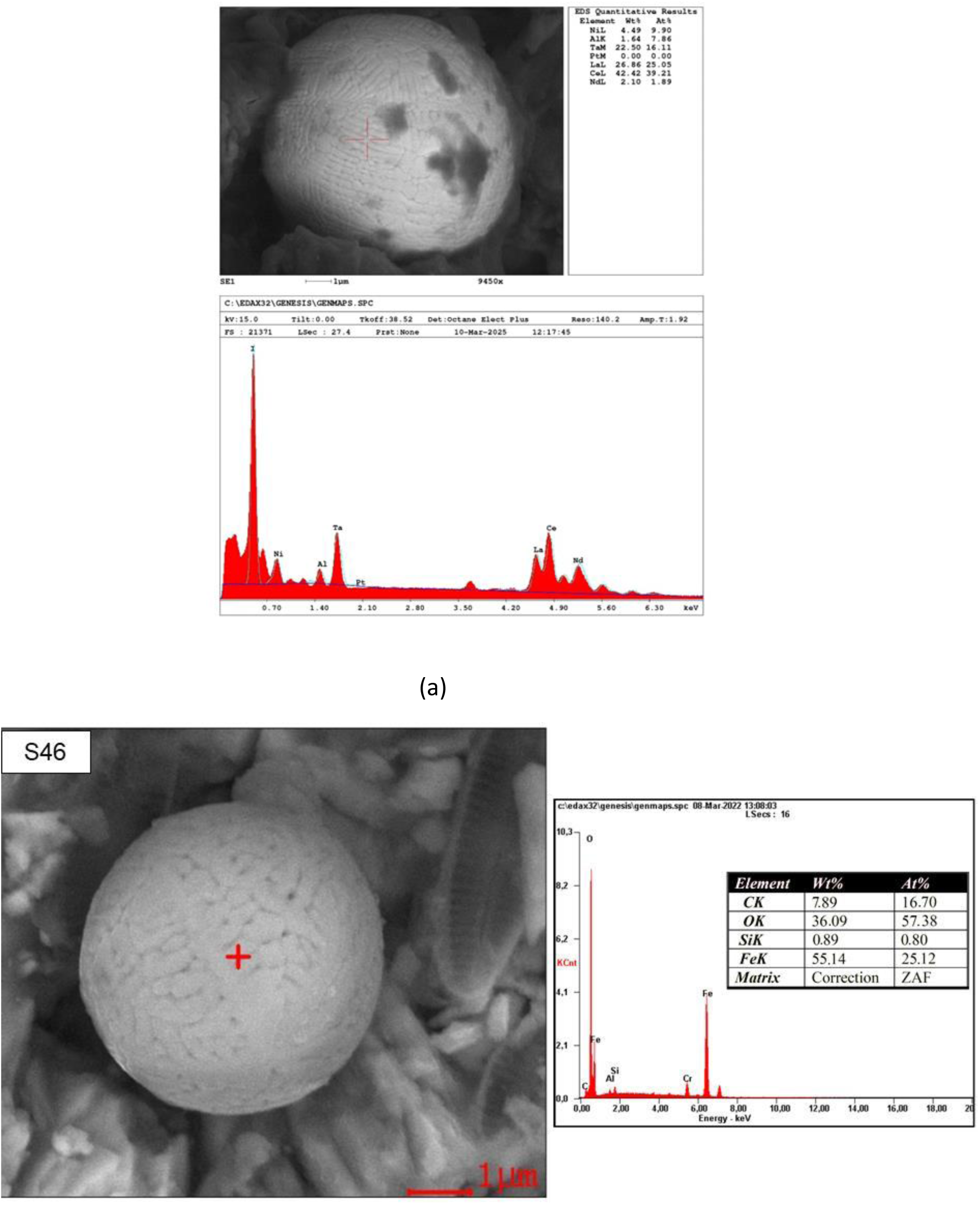
Semi-quantitative elemental chemical analysis of brain-like microspherules using a neighborhood electron microscope (SEM/EDS): a) Microspherule in the rock and b) Another microspherule in the rock, performed previously. Note the very different compositions of the two microspherules.

**Figure 25.**
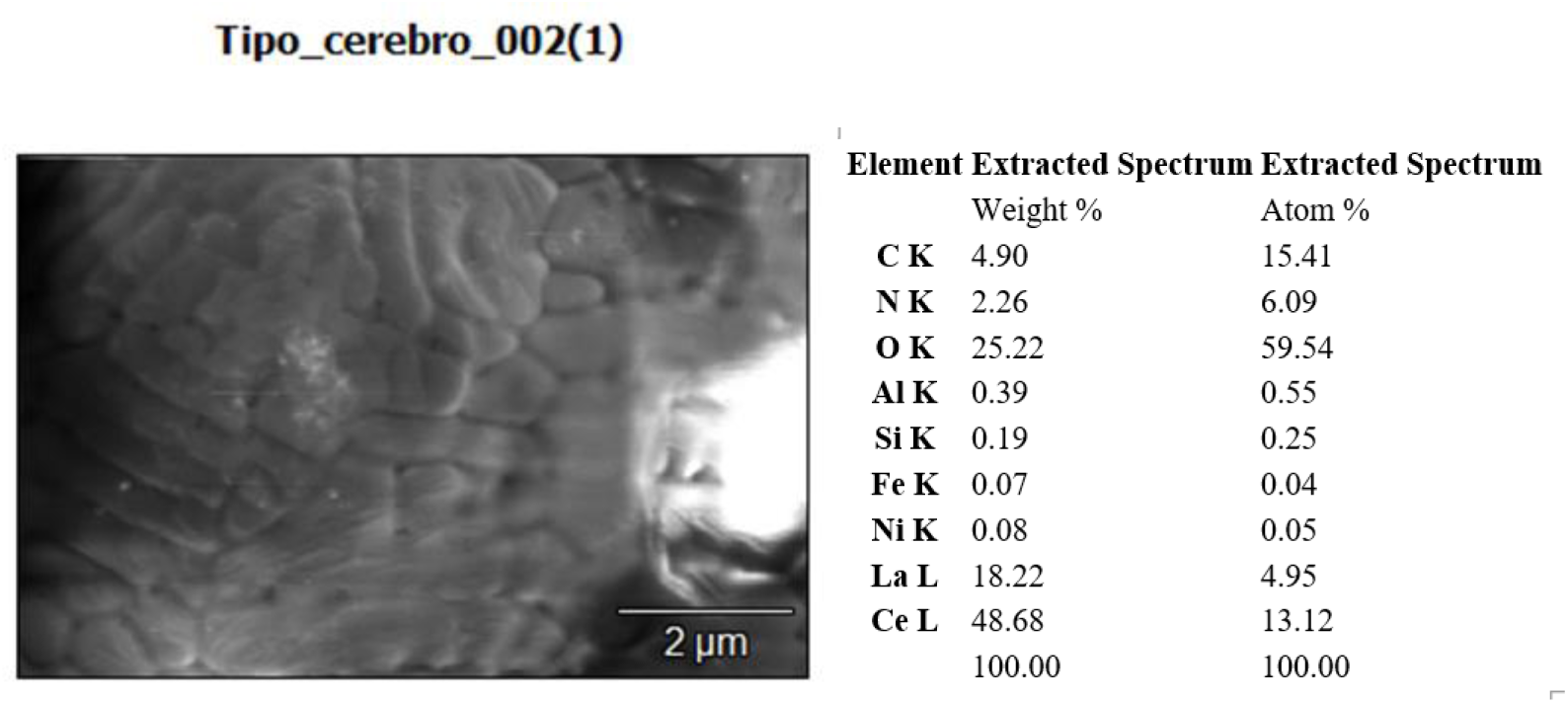
Dual beam (Backscattered) SEM EDS semi-quantitative elemental chemical analysis of brain-like microspherules: a) Microspherule in the analysis PIN. Note the similarity of their composition.

**Figure 26.**
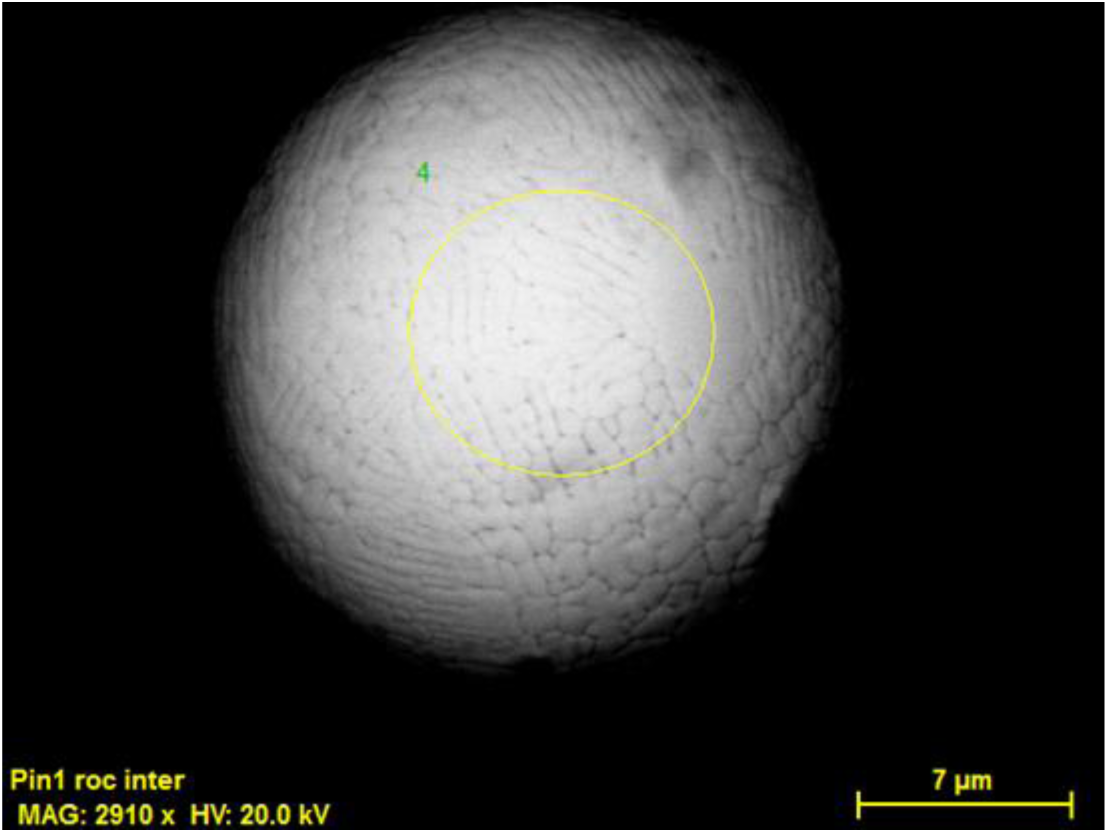
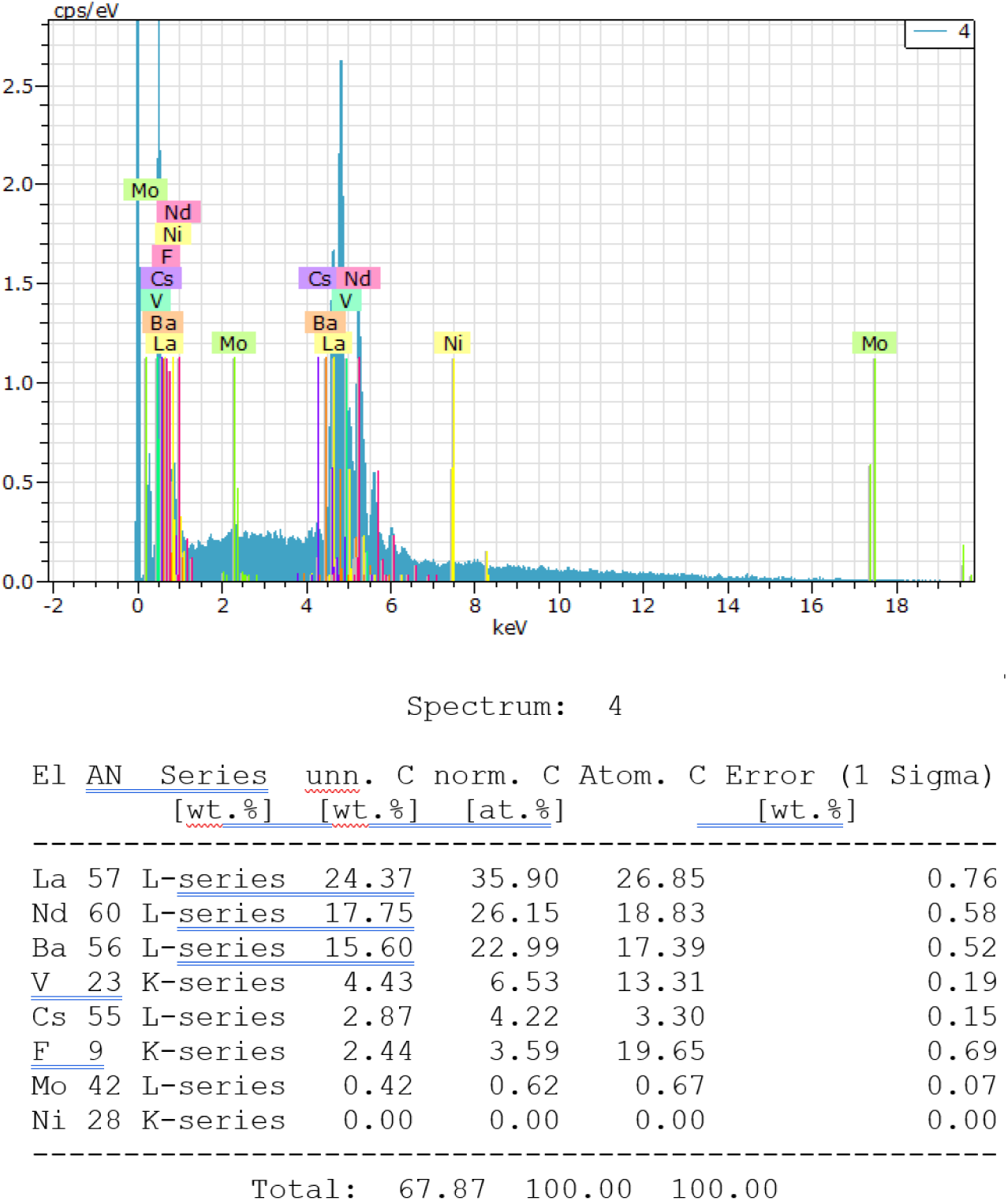
SEM EDS (Backscattered) semi-quantitative elemental chemical analysis of the brain-like microsphere on the graphene pin. Note the change in its composition.

Subsequently, another elemental chemical analysis was performed using a backscattered scanning electron microscope (SEM-EDS) at the National University of Colombia, Manizales, on the same brain-like microsphere in the graphene PIN. It was found that its composition had changed again (Figure 27). Samarium (Sm) was found, but no vanadium (V) was found.

**Figure 27.**
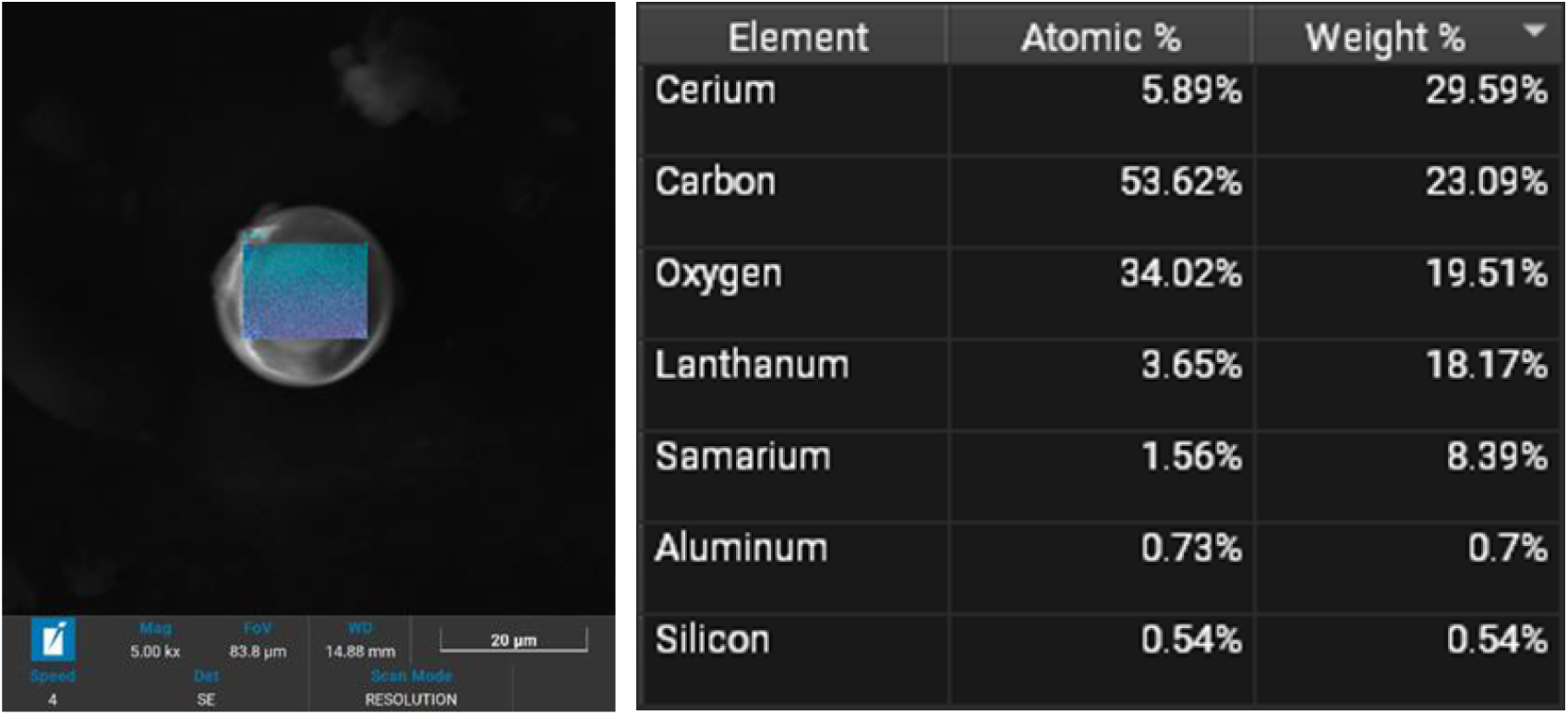
Backscattered SEM EDS (semi-quantitative elemental chemistry) analysis of the brain-like microsphere on the graphene pin. Note the change in its composition (Analysis performed at the National University of Colombia – Manizales campus).

Text taken from the document “Discovery of Spherules of Probable Extrasolar Composition at the Pacific Ocean Site CNEOS 2014-01-08 (IM1) Bolide, 2023:”

“We conducted an extensive towed magnetic sled survey during the period of June 14–28, 2023, over the seafloor approximately 85 km north of Manus Island, Papua New Guinea, and found approximately 700 spherules measuring 0.05–1.3 millimeters in diameter in our samples, of which 57 have been analyzed so far.

Approximately 0.26 km2 of seafloor was sampled in this survey, centered around the calculated trajectory of the CNEOS 2014-01-08 (IM1) bolide, with control areas to the north and south of that trajectory. The spherules, significantly concentrated along the trajectory, The meteorite’s mass spectrometry data are based on the expected abundance of the meteorite, from seafloor depths ranging from 1.5 to 2.2 kilometers. Mass spectrometry of 47 spherules near the high-yield regions along IM1’s trajectory reveals a clear extrasolar abundance pattern for five of them, while background spherules have abundances consistent with a solar system origin. The single spherules show an excess of Be, La, and U, up to three orders of magnitude relative to the solar system standard of CI chondrites. These previously unseen “BeLaU”-type spherules also have very low refractory siderophile elements such as Re. Volatile elements, such as Mn, Zn, and Pb, are depleted as expected due to evaporative losses during a meteor explosion. Furthermore, the mass-dependent variations in 57Fe/54Fe and 56Fe/Fe are also consistent with light loss due to isotope evaporation during the spherules’ journey through the atmosphere. The “BeLaU” abundance pattern is not found in control regions outside the trajectory of IM1 and does not match commonly fabricated alloys or natural meteorites in the Solar System. This evidence points to an association of “BeLaU”-type spherules with IM1, supporting their interstellar origin regardless of the high velocity and unusual material strength implied by the CNEOS data. We suggest that the “BeLaU” abundance pattern could originate from a highly differentiated magma ocean of an iron-core planet outside the Solar System or from more exotic sources. Figures 27 and 28, respectively.

**Figure 28.**
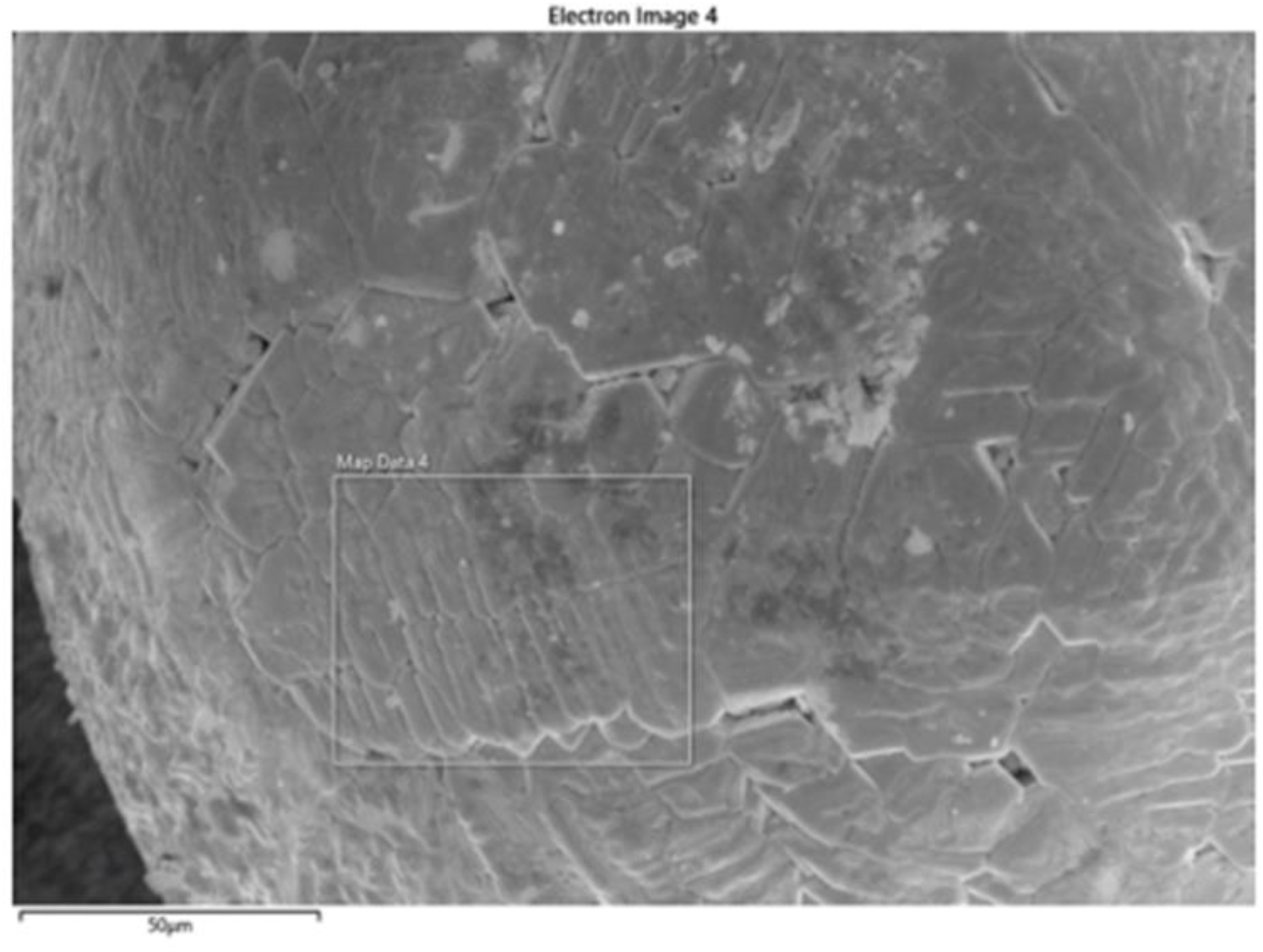
Figure C1. Spherule 14 from Experiment 8, showing an adult dendritic structure. Different regions display varying grain structure sizes, likely due to differences in thermal history (solidification rate, grain nucleation). Source: Discovery of Spherules of Likely Extrasolar Composition in the Pacific Ocean Site of the CNEOS 2014-01-08 (IM1) Bolide. 2023.

**Figure 29.**
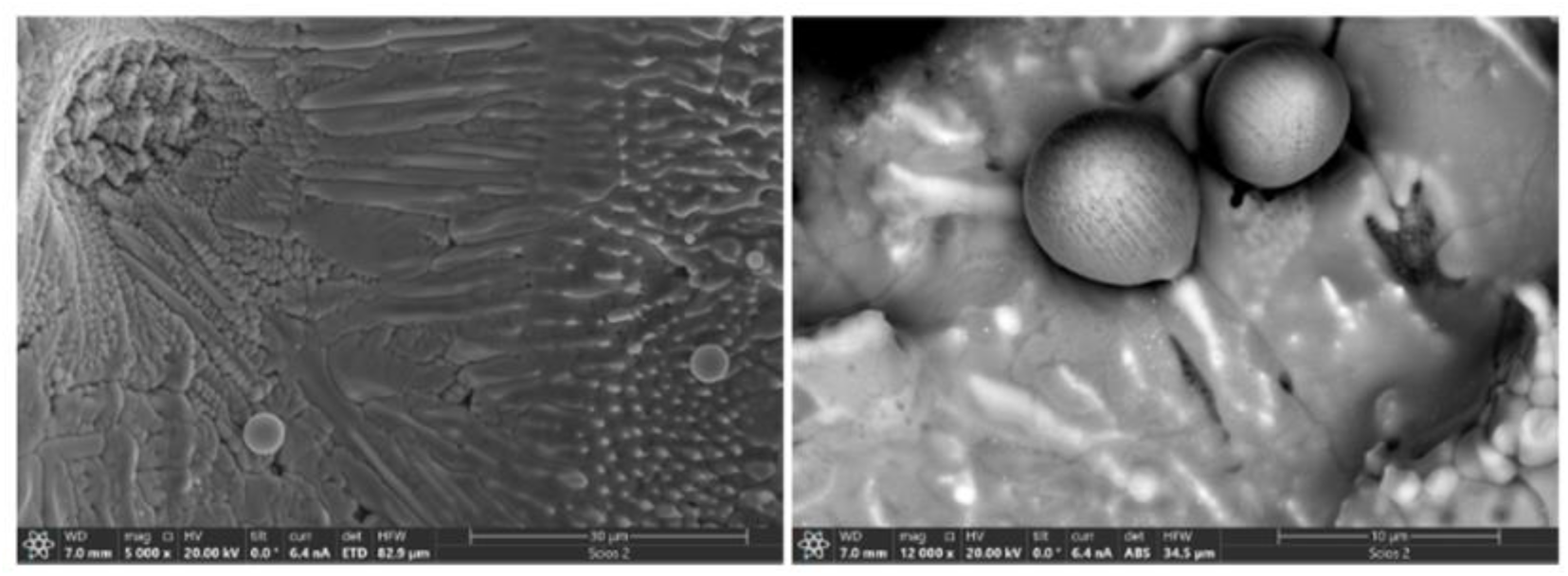
Figure C8. Spherule 4 from run 8 along the path of IM1, showing the dendritic structure. Note that several small, finely structured spherical particles are embedded in the core of larger particles. Source: Discovery of Spherules of Likely Extrasolar Composition in the Pacific Ocean Site of the CNEOS 2014-01-08 (IM1) Bolide. 2023.

Figures 28 and 29 show the similarity to the brain-like microspheres found in the rock.

**Table 1:** Collection of micrometeorites. Source:

| Deposits | Age | Collection Technique | No. Examined | Type | Size Range (µm) | Reference |
| --- | --- | --- | --- | --- | --- | --- |
| *Sediments* |  |  |  |  |  |  |
| Deep sea | ? | cores | 100s | S & I | 60–500 | Murray and Renard, 1883, 1891 |
| Deep sea | ? | magnetic rake | >300 | S & I | 100–500 | Brunn et al., 1955 |
| Deep sea | <7 Ma | core from top 3 m | 732 | S & I | 10–230 | Laevastu and Mellis, 1955 |
| Deep sea | <200 ka | magnetic sieve | 4413 | ? | 30–250 | Pettersson and Fredriksson, 1958 |
| Deep sea | ? | 750 kg sieved, mag. sep. | 1200 | I | 149–351 | Millard and Finkelman, 1970 |
| Deep sea | 0–100,000 yr(?) | magnetic rake | 100s | S, G & I | ≤5000 | Brownlee et al., 1979 |
| Deep sea | 0–500,000 yr | mag. sep. box core | >700 | S, G & I | 100–000 | Blanchard et al., 1980 |
| Deep sea | 0–700,000 yr | 1 m clam shell sample | 935 | S & I | 149–750 | Murrell et al., 1980 |
| Deep sea | 0–350,000 yr | 0–35 cm core, mag. sep. | 258 | S, G & I | 50–500 | Kyte, 1983 |
| Desert sand | ? | magnetic collector | 32 | I | 30–300 | Fredriksson and Gowdy, 1963 |
| Beach sand | recent–1.6 Ma | hand magnet | ? | I | 80–650 | Marvin and Einaudi, 1967 |
| *Sedimentary rocks* |  |  |  |  |  |  |
| Hardgrounds | 145–185 Ma | crush and dissolve | ? | I | 100–300 | Czajkowski et al., 1983 |
| Hardgrounds | 180 Ma | ? | 12 | I | ? | Jehanno et al., 1988 |
| Claystones | recent–500 Ma | dissolution, magnetic sep. | ? | ? | <40 | Crozier, 1960 |
| Carbonates | 30–40 Ma | dissolution, magnetic sep. | 28 | S & I | >100 | Taylor and Brownlee, 1991 |
| Salt deposits | ~250 & ~400 Ma | dissolution, magnetic sep. | 243 | ? | <40 | Mutch, 1966 |
| Sandstones | 1.4 Ga | mineral separation | 4 | S | 60–125 | Deutsch et al., 1998 |
| *Polar ice and sediment* |  |  |  |  |  |  |
| *Greenland* |  |  |  |  |  |  |
| Cryoconite | ? | heavy liq. separation | ? | S? | 100–200 | Wulfing, 1890 |
| Cryoconite | 0–3000 yr | suction, filter and pick | ~3500 | all | 50–300 | Maurette et al., 1987 |
| Cryoconite | <2 ka | suction, filter and pick | >100 | all | 100–1000 | Maurette et al., 1986 |
| *Antarctica* |  |  |  |  |  |  |
| Eolian deposits | <2.5 Ma(?) | sieved and hand picked | 840 | S | 125–500 | Hagen et al., 1990 |
| Eolian deposits | 0–100,000 yr(?) | wet sieved and hand picked | >100 | S | 64–1000 | Harvey and Maurette, 1991 |
| Ice cores | 1800–1961 AD | melt and filter | ? | I | 15–180 | Thiel and Schmidt, 1961 |
| Ice cores | 2300–100 BC | melt and filter | 5 | S, G, V | 50–160 | Yiou and Raisbeck, 1987 |
| Ice | ? | melt and filter | 76 | all | 50–400 | Maurette et al., 1991 |
| Snow and ice | 1100–1500 AD | suction bottom and filter | 1600 | all | 50–800 | Taylor et al., 1998 |

Micrometeorites are classified according to the intensity with which they have been heated (cosmic spherules or fully melted, partially melted, and unmelted) and by their composition (chondritic versus nonchondritic). Figure 30 shows examples of some of the main types.

**Figure 30.**
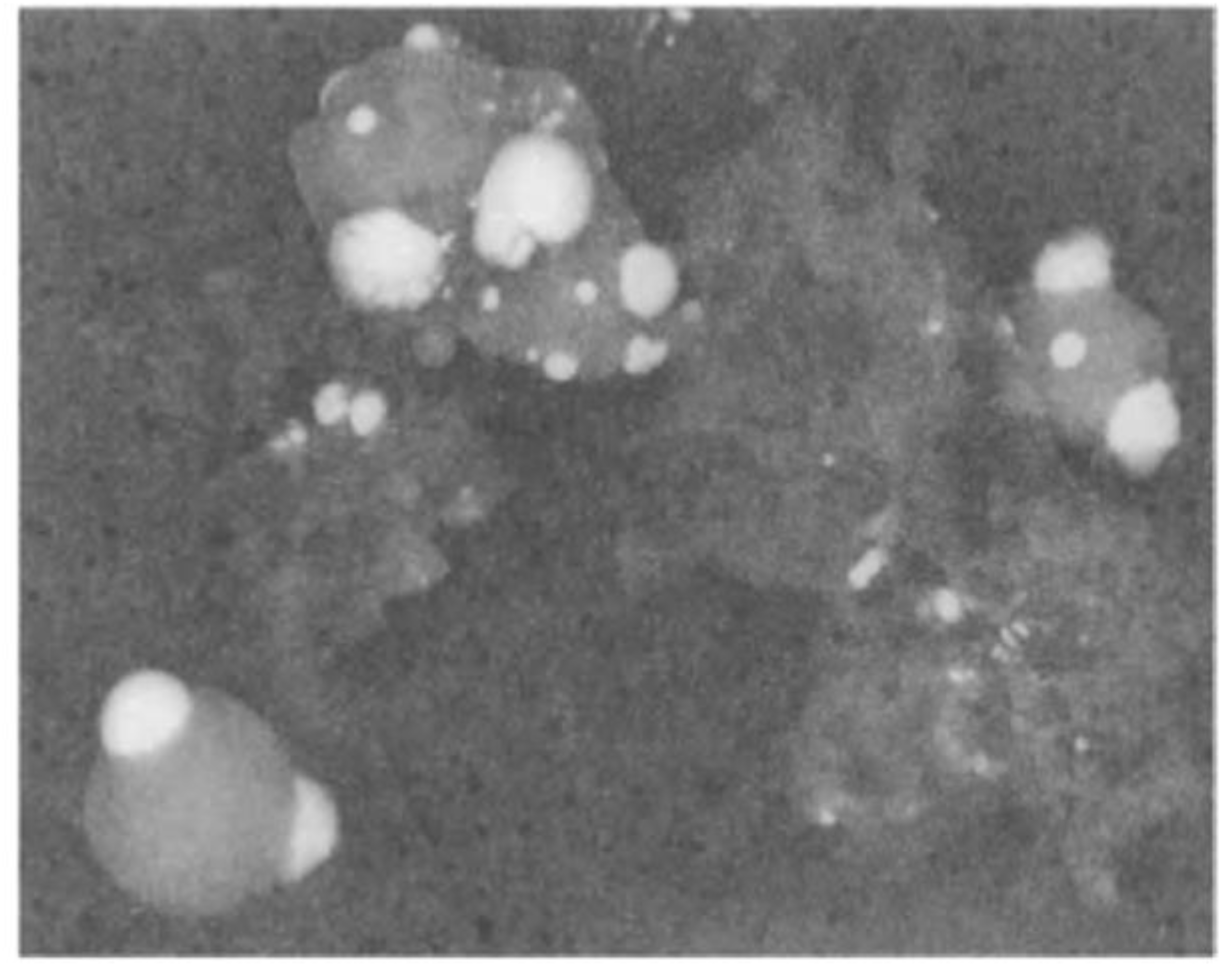
From Figure 3. Backscattered SEM image of a single interplanetary dust particle that melted during atmospheric entry to form silicate spherules coated with metal beads (bright in the image). The irregular components are carbon that survived atmospheric entry and the more refractory component of the extraterrestrial matter accreted from space. Source: Accretion of Extraterrestrial Matter Throughout Earth’s History Edited by Bernhard Peucker-Ehrenbrink Woods Hole Oceanographic Institution Woods Hole, Massachusetts and Birger Schmitz Earth Sciences Center, University of Gothenburg Gothenburg, Sweden 2001 Springer Science+Business Media New York. Note the similarity to the planet-like microspheres (Figure 28 (S30)

Iron spherules (type I) are composed of magnetite crystals intertwined with interstitial wiristite (Fig. IA in Figure 30). They may be the oxidized and melted version of the rare-metal grains described by Maurette et al. (1987), or possibly immiscible metallic phases that separated from other micrometeorites by buoyancy during atmospheric entry.

Type G spherules have a high iron content and contain magnetite dendrites in a silicate glass matrix (Fig. IB in Figure 30). Their low Mn content suggests that they are not chondritic (Brownlee et al., 1997).

The stony spherules (type S) have chondritic compositions and show a range of heating effects from fully melted to partially melted with relict grains and scoriaceous particles. These include (i) glass spherules, composed of mafic glass that has been fully melted, devolatilized, and rapidly quenched (Fig. IC in Figure 30), (ii) cryptocrystalline spherules that have no visible magnetite or olivine crystals but show structure under cross-polarization (Fig. 1D in Figure 30), and (iii) barred olivine spherules that have also been fully melted but have cooled less rapidly, allowing olivine and magnetite crystals to form (Fig. IE in Figure 30). Porphyritic spherules have equidimensional olivine crystals and magnetite crystals in interstitial glass. They sometimes feature commonly zoned olivine grains with forsterite cores and fayalitic rims (Fig. IF in Figure 30). Spherules containing relict grains were not completely melted and did not retain precursor minerals, usually relict olivine, metal, or sulfide grains (Fig. IG in Figure 30). Scoriaceous particles are partially melted vesicular spherules containing relict olivine, metal, or sulfide grains in glass (Fig. 1H in Figure 30).

Unmelted, fine-grained micrometeorites have dark, tightly packed phyllosilicate matrices containing metal, sulfide, and silicate grains. These have chondritic compositions, and magnetite rims surround the particles and most of the vesicles (Fig. II). Unmelted coarse-grained micrometeorites (Fig. 1J in Figure 30) contain crystals of pyroxene, olivine, and plagioclase in a glassy matrix. These particles do not have chondritic compositions, and cosmic spherules with a composition of pyroxene and olivine (Brownlee et al., 1993; Taylor et al., 2000) can form when coarse-grained monomineral micrometeorites are melted. Source:

### Planet-like Microchondrules

In Cluster 2 (M2), there were 29 planet-like, smooth, and spherical microchondrules, some with lobes, bumps, spots, and scales (Fig. 31), and they were predominantly small. At least 10 of them contained C, O, Si, Fe, Al, Ca, Ni, La, and Ce. Other elements, such as N, Mg, and F, were present in at least five of them. This group had two microspheres that, while not smooth, had similar attributes, such as being spherical and having scales. Both had high iron contents; one of them was practically pure iron (92%), and the other was composed of C, N, O, and Fe (47%) (Fig. 32). Furthermore, a microsphere was identified within the group that could be in the process of subdivision, although the section that appeared to be separating differed in elemental composition with high Fe content and reduced La and Ce values (Fig. 33a). Figure 33b shows a possible budding occurring in one of these microspheres, showing that a parent microsphere has different chemical elements than the offspring, while Figure 33c shows a possible hatching.

**Figure 31.**
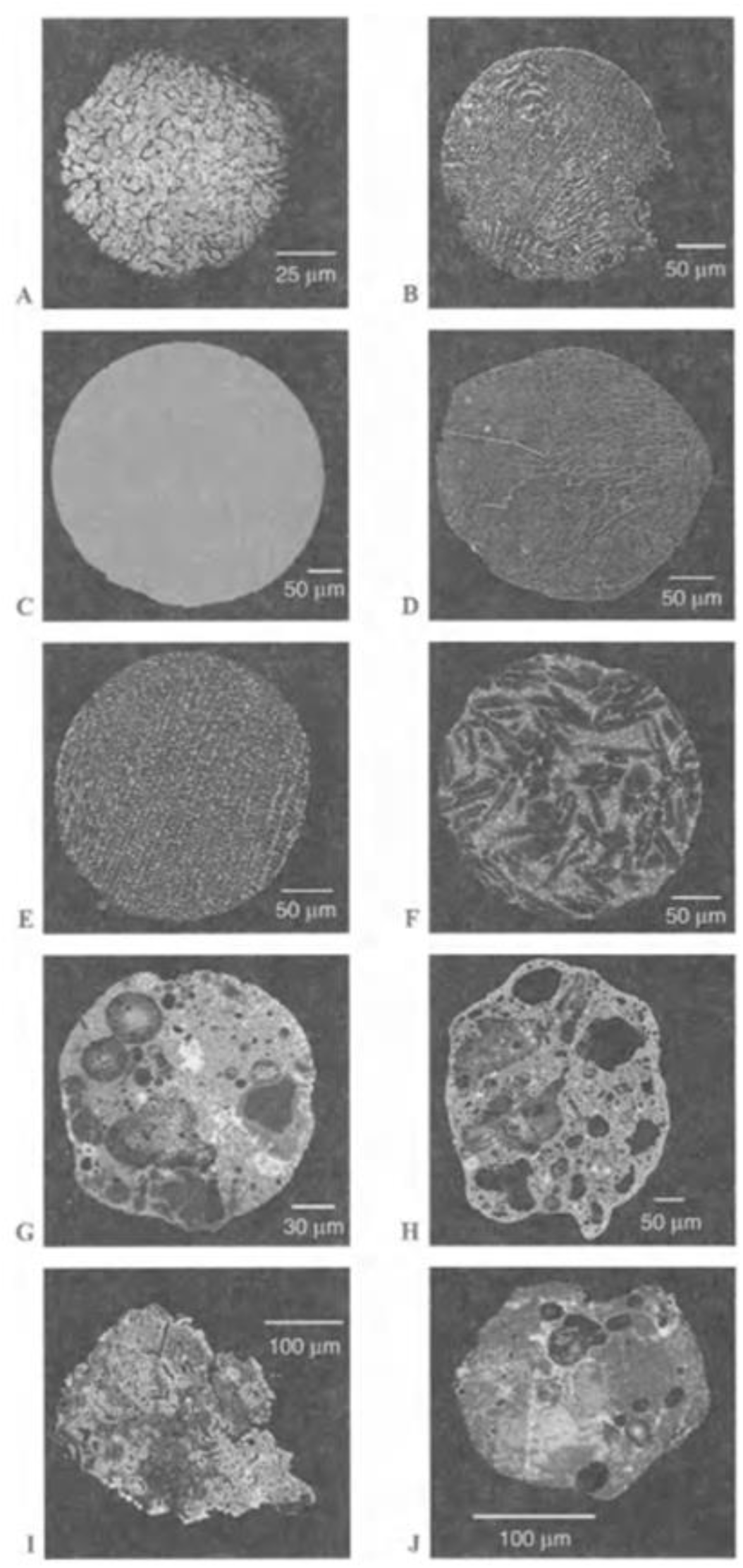
Examples of the main micrometeorite types. (A) Iron spherule (type I) composed of magnetite crystals with interstitial wiristite, some of which have disappeared. (B) Type G spherule with magnetite dendrites (bright phase) in a glass matrix. (C) Glass spherule composed of mafic glass. (D) Cryptocrystalline spherule without visible magnetite or olivine crystals, but showing flow features. (E) Barred olivine spherule with lathe-shaped olivine crystals (dark phase) and small magnetite crystals (bright phase) in interstitial glass. (F) Porphyritic spherule with equidimensional olivine crystals and magnetite crystals in interstitial glass. The dark areas in some of the olivines are of forsteritic composition. (G) Spherule containing relict grains of olivine, metal, and sulfide, indicating that the particle has not completely melted. (H) Scoriaceous particle with relict grains of olivine, metal, and sulfide in highly vesicular glass. (I) Unmelted, fine-grained micrometeorite showing a dark, tightly packed phyllosilicate matrix containing metal, sulfide, and silicate grains. Magnetite rims surround the particle and most of the vesicles. The lighter-colored regions have been intensely heated. (J) Unmelted, coarse-grained micrometeorite composed of olivine (light gray), orthopyroxene (gray), and chromite (bright phase) in a glass matrix. Image provided by Dr. Gero Kurat. (Panels A, E, G, and I are in Taylor et al., 2000, reproduced with permission.)

**Figure 32:**
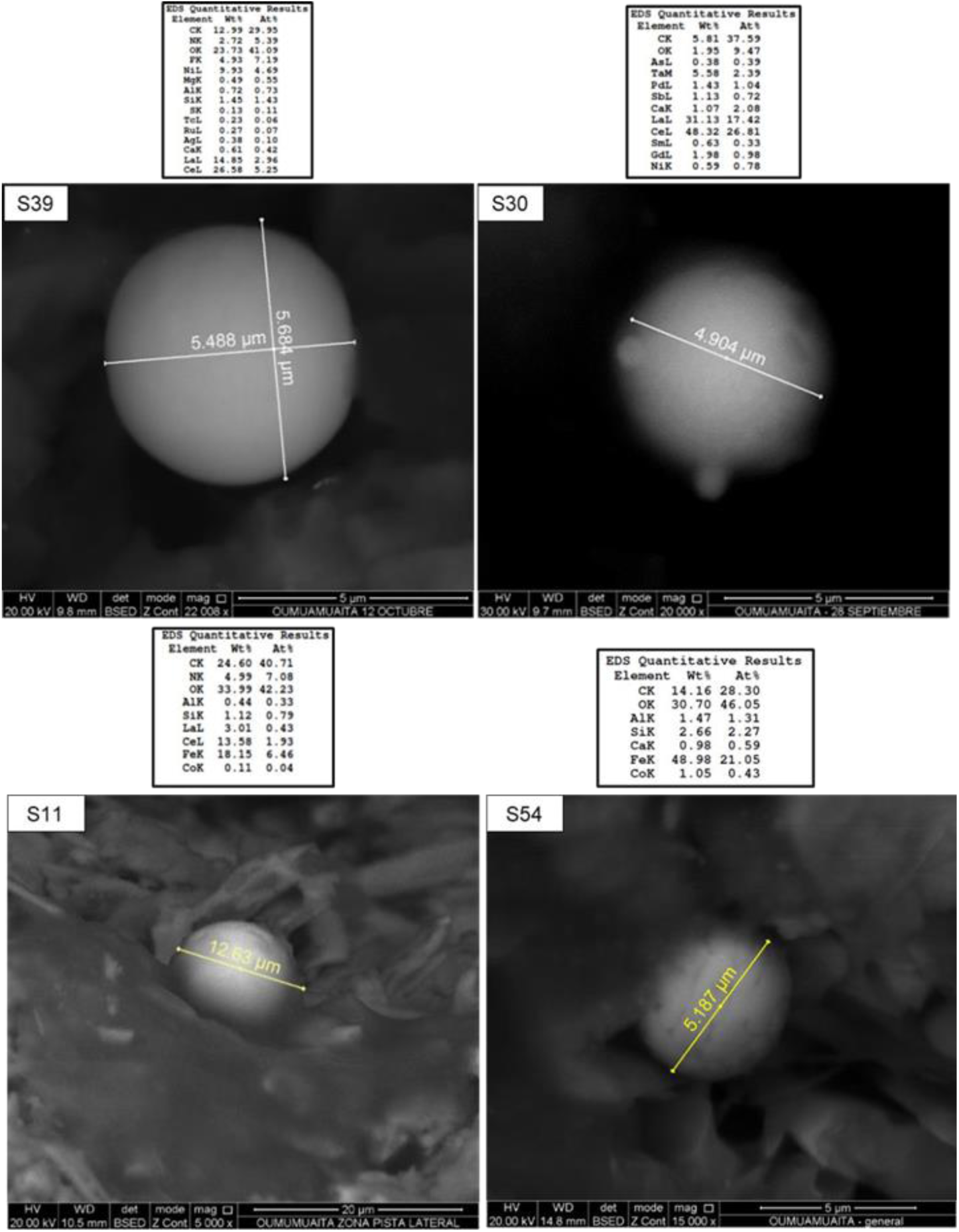
Smooth microspheres with bumps, lobes, scales, or spots, composed primarily of carbon, oxygen, lanthanum, and cerium or iron. Semiquantitative chemical analysis performed using a scanning electron microscope (SEM-EDS).

**Figure 33:**
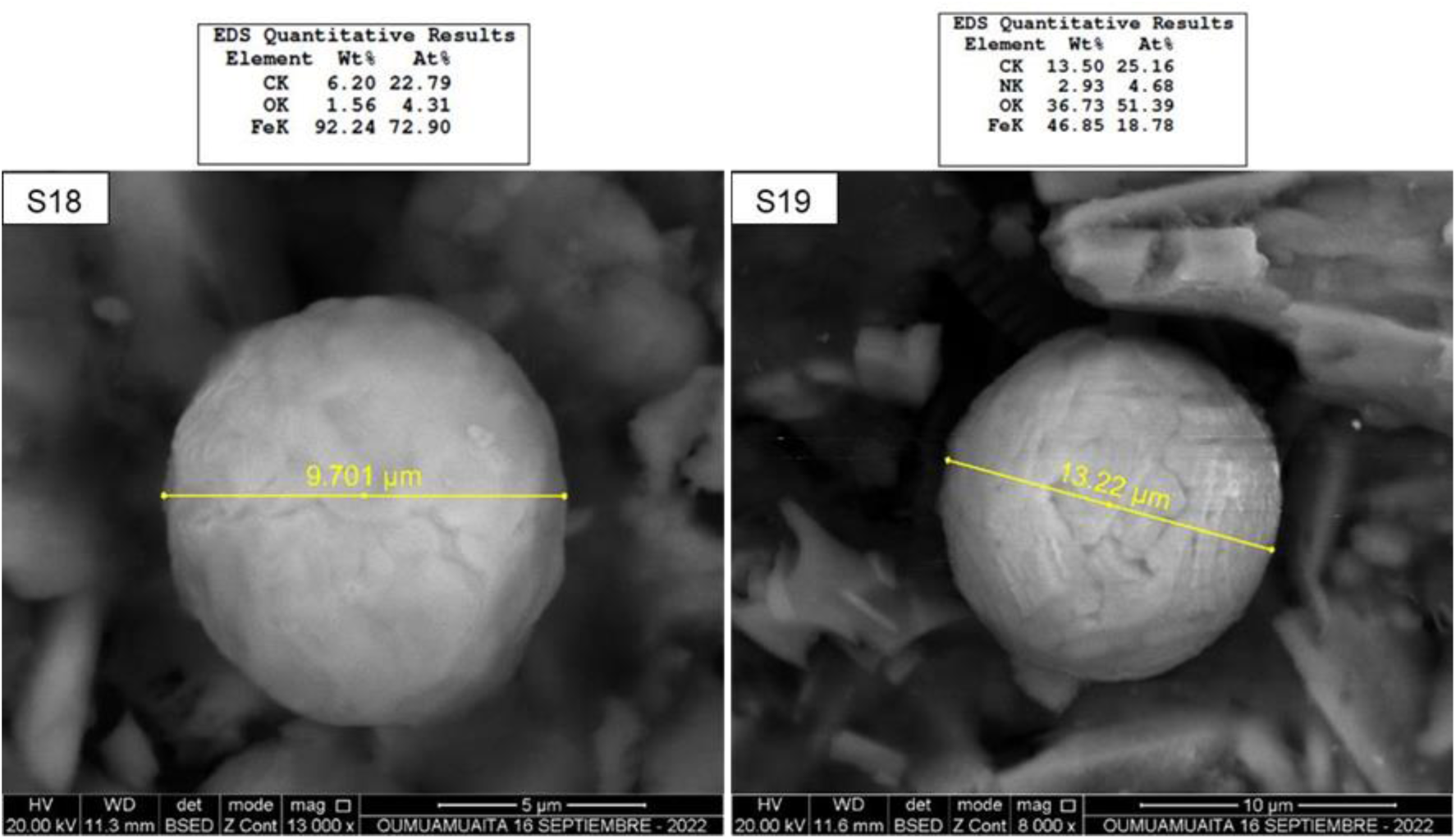
Microspheres with rough, spherical morphology and with the presence of scales composed of carbon, oxygen and iron. Semiquantitative chemical analysis carried out with a scanning electron microscope (SEM – EDS).

### Lemon-like Microchondrules

In Group 3 (M3), there were ten mainly ovoid, lemon-like microchondrules, small in size, and distributed between granules and smooth (Fig. 34). Their characteristic elements included C, O, and Si in all of them, as well as Fe, N, Mg, Al, Ca, and Na in at least three of them. One microsphere with a lemon-like appearance stood out due to its morphology, which may be a developmental stage of the granular stage, and was the largest of the group (Fig. 35).

**Figure 34.**
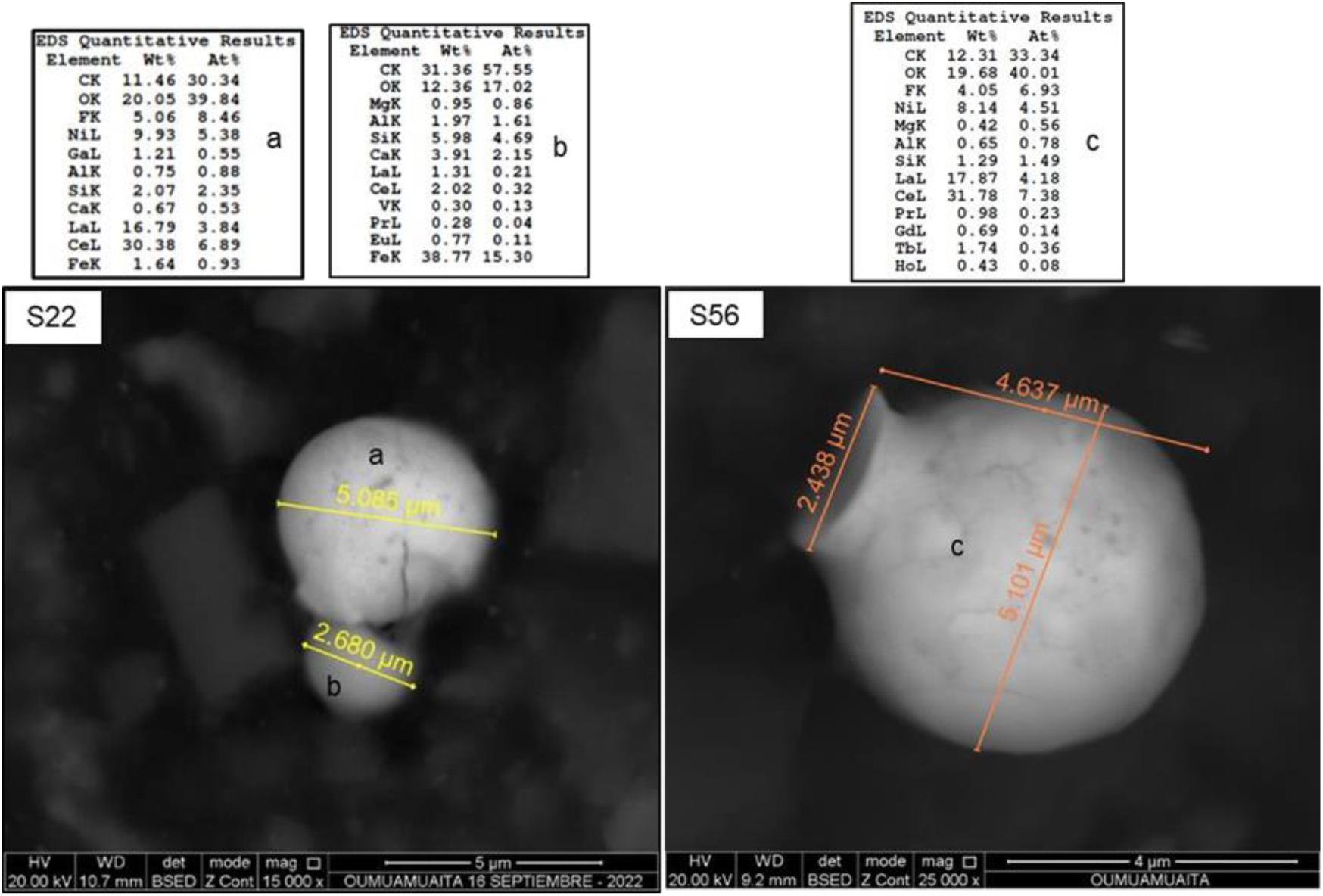
Smooth, mottled microspheres undergoing apparent subdivision and possibly hatching, composed primarily of carbon, oxygen, lanthanum, and cerium. Semiquantitative chemical analysis performed using a scanning electron microscope (SEM) – EDS.

**Figure 35:**
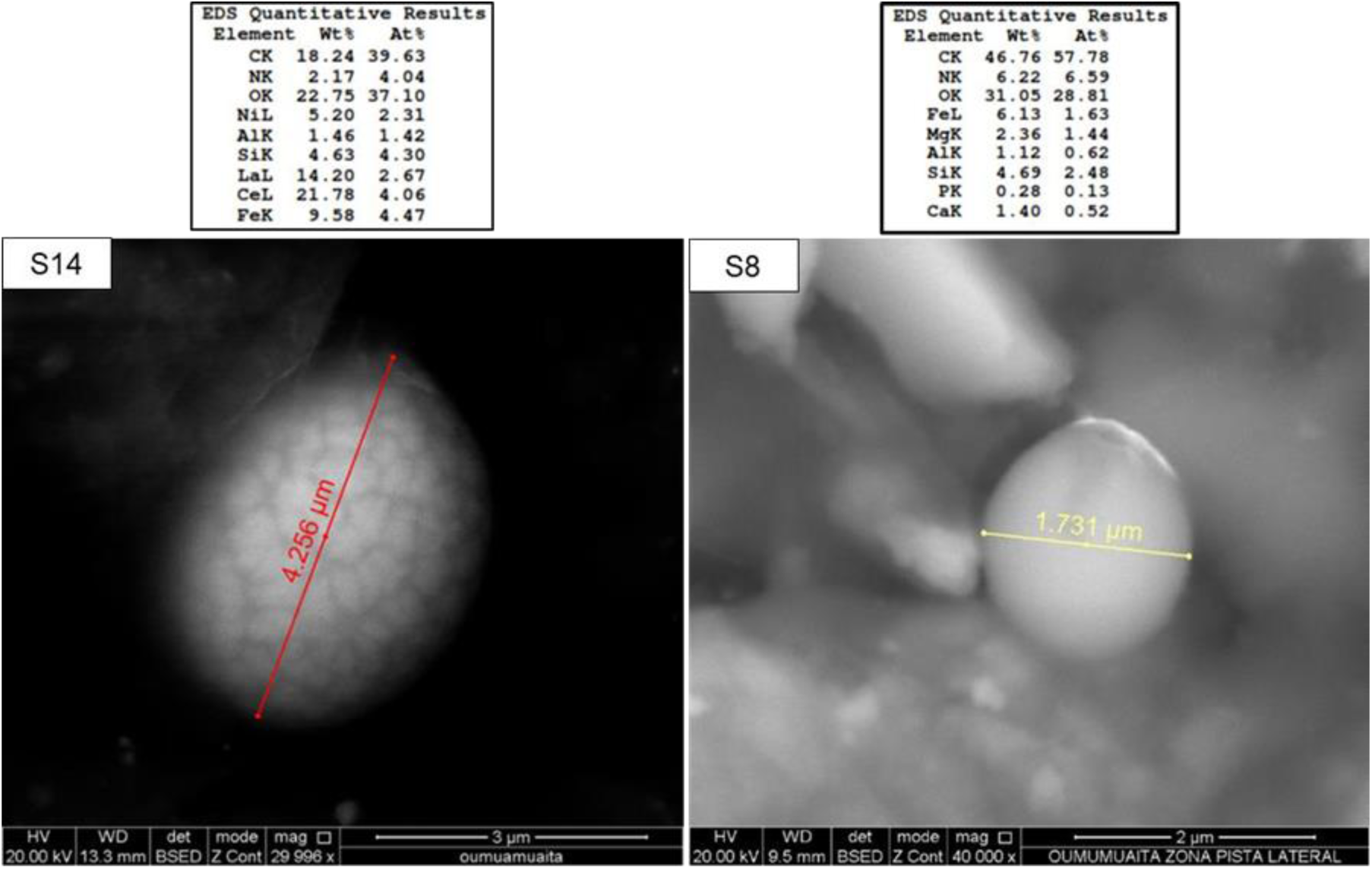
Microspheres with an ovoid morphology and composed mainly of carbon, oxygen and lanthanum and cerium or iron and silicon. Semiquantitative chemical analysis carried out with a scanning electron microscope (SEM – EDS).

**Fig. 35:**
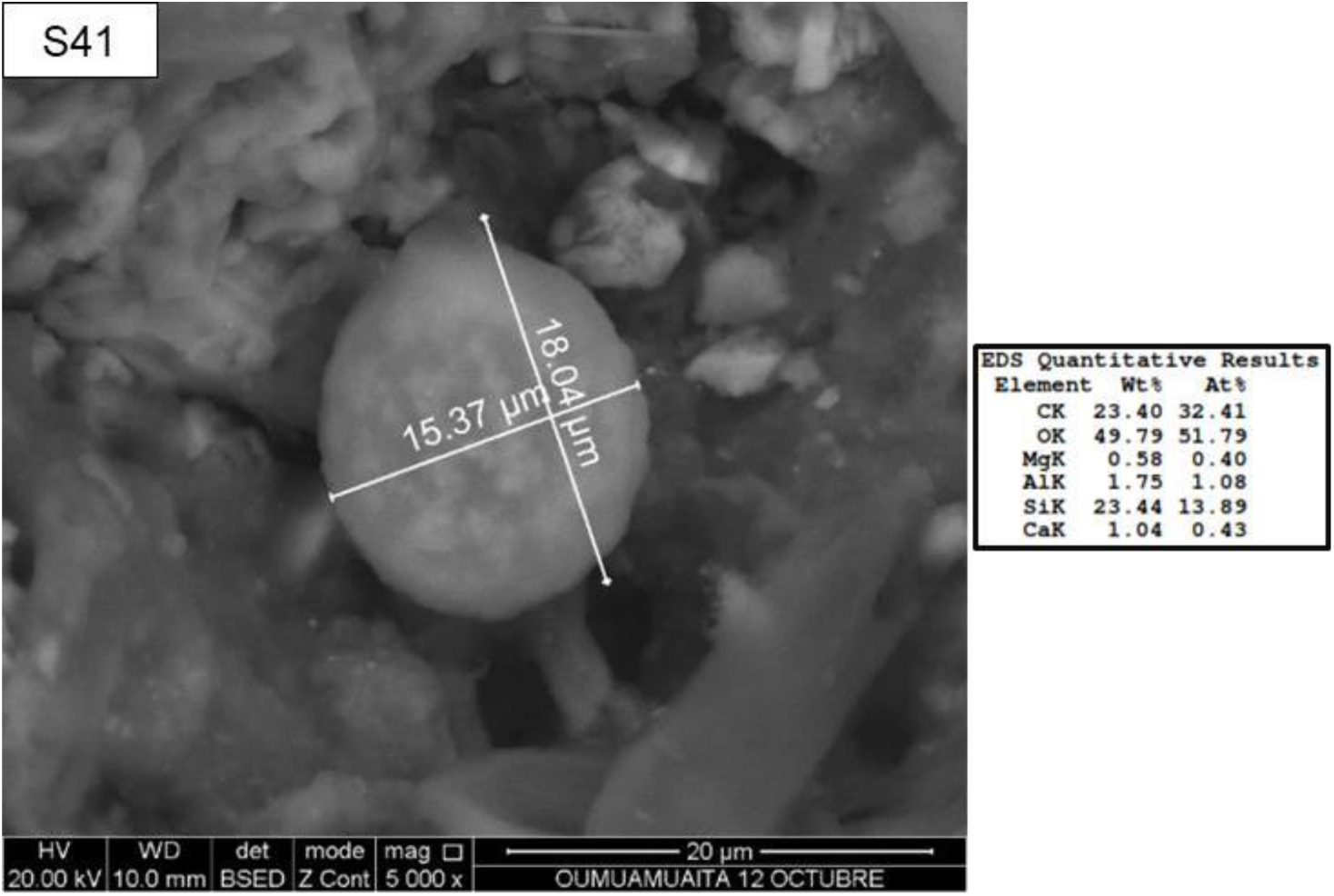
Ovoid microsphere with mottled morphology and composed primarily of carbon, oxygen, and silicon. Semiquantitative chemical analysis performed using a scanning electron microscope (SEM-EDS).

### Granular Microchondrules

Group 4 (M4) included thirteen small to medium-sized microchondrules with a granular and spherical appearance (Fig. 36). Some had bumps, spots, and scales. The elements identified were C, O, and Si in 11 of them, as well as Fe, Al, Ni, Ca, La, Ce, and Ni in at least seven of them.

**Figure 36:**
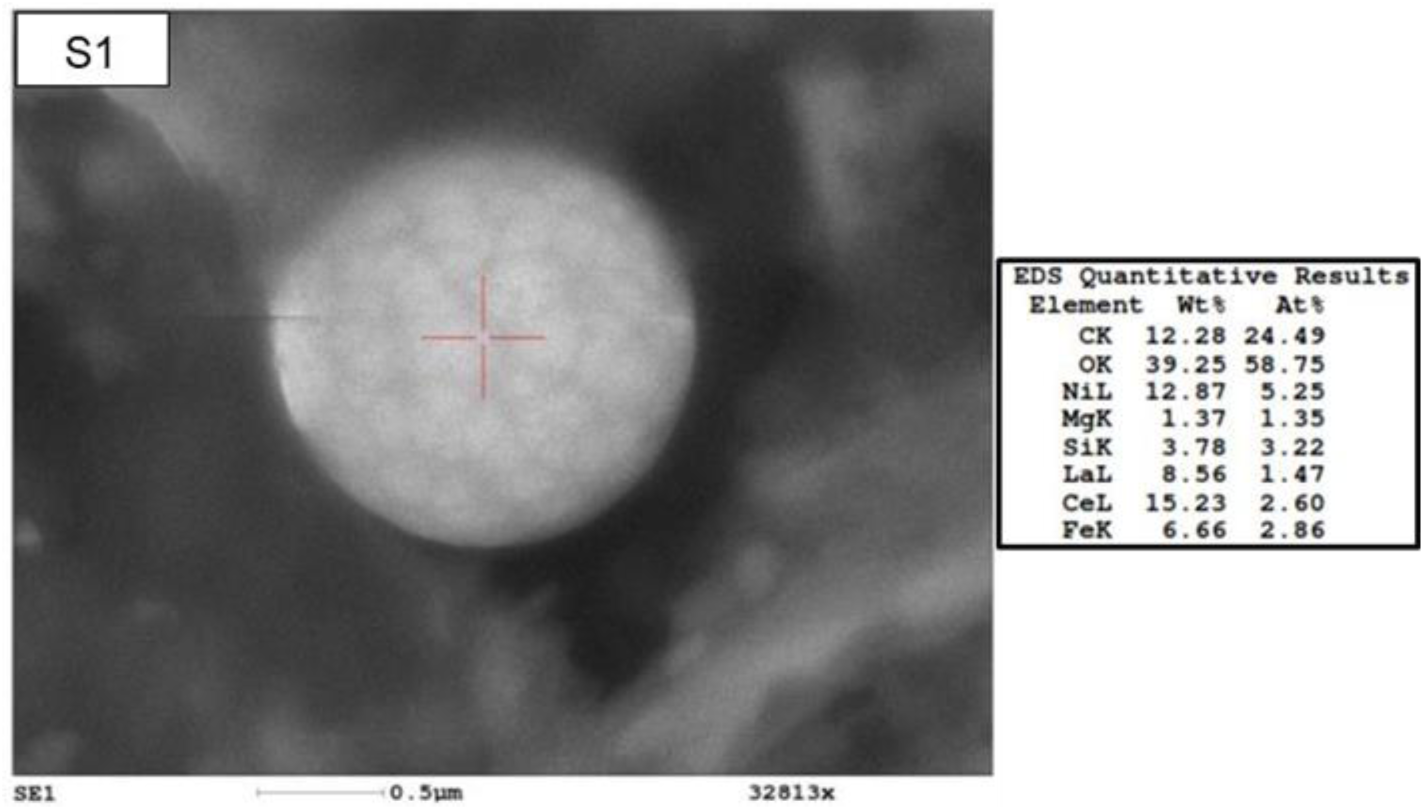
Microsphere with a granular morphology and composed primarily of carbon, oxygen, cerium, lanthanum, and nickel. Semiquantitative chemical analysis performed using a scanning electron microscope (SEM) – EDS.

### What are these microchondrules or microspheres?

Based on the above, a set of potential entities or microchondrules observed by scanning electron microscopy (SEM) were recorded and chemically analyzed (quantitative analysis). This showed that they are not inorganic particles; these microchondrules are organic and composed of carbon, oxygen, nitrogen, silicon, aluminum, iron, calcium, lanthanum, cerium, titanium, vanadium, and nickel, among others. Therefore, they may have an organic composition.

These organic microchondrules, both in size and morphology, are distinct from known microorganisms and are not associated with Fe(Mn) oxide spherules as described by Kos et al. (2022), pollen, grass seeds, or other terrestrial organisms such as bacteria, or with the Kingdom Chromista (photosynthetic protists and related protists) or with the Kingdom Protozoa (primitive protozoa and most slime molds).

Comparing these results with analyses performed on samples brought from the stratosphere at altitudes of 23–27 km (Wainwright, 2015), the similarities are striking, and these results are even more surprising because such a wide variety of these different microchondrules were found on the surface of such a small rock. Based on the above, the possibility arises that these microchondrules may have arrived from space and therefore do not have a terrestrial origin.

On the other hand, contrary to the explanations for the origin of the spheres reported in samples from the stratosphere at altitudes of 23-27 km, which indicate a provenance from space (Wainwright, 2015), it is possible to affirm that microchondrules with a spherical morphology coexisted on the surface of the recovered rock and were not the only microchondrules that could be found in this material. However, microspheres were among the most representative, and due to their characteristics of mixing organic materials with metals and other elements unusual in entities known on Earth, they allow us to consider a possible non-terrestrial origin.

There is no conclusive evidence that the rock is of meteoritic origin, and in principle, it should be terrestrial. On the contrary, its composition, derived from the presence of a large number of chemical elements, could indicate a non-terrestrial origin. In this case, the length of time the rock remained submerged in water before being recovered is unknown. Therefore, it is possible that some of the observed materials are terrestrial products. However, in any case, given that general analyses of the rock indicate that it is of terrestrial origin, and those produced by SEM, which show similarity to microspheres recovered from the stratosphere, indicate a possible non-terrestrial origin, it is necessary to consider the existence of new organic microchondrules that have not yet been scientifically identified and described. Most of them have been observed for the first time, and at least one of them exhibits a potential biological entity due to a budding process.

## Conclusions

This study contributes to the understanding of new life forms based on chemistry distinct from that known on our planet, which can aid in the development of new future research in various fields of science, such as biology, chemistry, geology, and genetics, among others.

Through SEM and EDS analysis, it was determined that the sum of all microchondrules with spherical morphology contained up to 48 elements, the most important of which were C, O, N, Si, Ti, V, Ni, La, and Ce. They were not associated with Fe (Mn) oxide spherules, pollen, grass seeds, or other terrestrial organisms such as bacteria, the Kingdom Chromista (photosynthetic protists and related protists), or the Kingdom Protozoa (primitive protozoa and most slime molds).

The observed microchondrules were classified according to their composition and morphology into three and four groups, respectively, demonstrating that they share some characteristics, such as the presence of carbon and oxygen, and other very common elements such as Si, Al, Fe, and Ca. Some microspheres showed high contents of rare earth elements, especially lanthanum and cerium, and very high contents of metallic elements not known in organic structures, such as Ti, V, Au, Pt, Ba, and Ni, Sm, La; Cs, Ce, Nd, Nb, among others.

Comparing these results with the analyses performed on stratospheric samples, the similarities are striking, and the results presented are even more so, since a large number of these different microchondrules are found in such a small rock.

From a morphological perspective, at least four major categories were identified, dominated by brain-like, granular, and smooth morphologies, and both spherical and ovoid types. The different sizes found in the same morphological type may be evidence of a process of growth and development of microchondrules, combined with a budding process, which makes it possible to observe biological entities.

Based on the above, and given that these microchondrules are of organic composition, according to the panspermia hypothesis, the possibility arises that this rock arrived from space and, therefore, microchondrules are possible biological entities that do not have a terrestrial origin.

To strengthen the conclusions and align the findings with established scientific practices, a proposal for **new taxonomic nomenclature is essential**. Given the unique characteristics described in the article, the following nomenclature is proposed, keeping in mind that formal **taxonomic classification** would require further research and validation:

### Proposed Nomenclature

Kingdom: Astromicrobiota: (signifying a potential origin beyond Earth)

Phylum: Sphaeromorphida: (referencing the predominant spherical morphology)

Class: Based on morphological distinctions, the microchondrules can be divided into classes such as:

1.- Cerebromorpha (for brain-like structures) 2.-Granulomorphea (for granular structures) 3.-Leiomorpha (for smooth structures)

4.- Ovomorpha (for ovoid structures)

Order: Further division based on elemental composition: 1.- Ferrochondrales (high iron content)

2.- Silicochondrales (high silicon content) 3.- Carbochondrales (high carbon content)

Family: Microchondrulaceae (general family designation) Genus: Microchondrulus (general genus designation)

Species: Species names would be based on specific distinguishing characteristics, such as the location of discovery or unique elemental markers.

Examples:

1.- Astromicrobiota sphaeromorphida cerebromorpha ferrochondrales microchondrulaceae

*Microchondrulus columbiensis* (brain-like, iron-rich, found in Colombia)

2.- Astromicrobiota sphaeromorphida granulomorphea silicochondrales microchondrulaceae *Microchondrulus granularis* (granular, silicon-rich, general)

Rationale:

The term “Astromicrobiota” reflects the hypothesis of an extraterrestrial origin, aligning with the panspermia implications discussed in the article’s introduction.

The use of established taxonomic ranks (Phylum, Class, Order, Family, Genus, Species) provides a framework for future research and comparison.

Morphological and elemental characteristics are used as the primary basis for classification, reflecting the key findings of the study.

Next Steps:

1. Further Research: In-depth genomic and proteomic analysis is required to determine if these entities contain unique genetic material.
2. Replication: Independent verification of these findings by other research teams is crucial.
3. Culturing: Attempts to culture these entities would provide invaluable insights into their biology.
4. Formal Classification: If further research supports a unique taxonomic position, a formal proposal should be submitted to the appropriate taxonomic authority.

By proposing this nomenclature, the study can move beyond mere observation and provide a structured framework for future exploration into the nature and origin of these intriguing microchondrules.

## Acknowledgments

This group of researchers wishes to acknowledge Dr. Marco António Ramírez Olvera and Dr. Margarita Reyes for their support in the analysis of LA ROCA at the Mexican Institute of Geology, UNAM, Scanning Electron Microscopy (SEM) Laboratory.

## Notes

### Competing Interest Statement

The authors have declared no competing interest.

## Bibliographic references

Badyukov, D.D., Raitala, J. Ablation spherules in the Sikhote Alin meteorite and their genesis. Petrology 20, 520–528 (2012). 10.1134/S086959111206001X

Bignami, L., Guaita, C., Pezzotta, F., & Zilioli, M. (2014). Micro-spherules near the Kamil crater. ResearchGate. https://www.researchgate.net/publication/281787526_Micro-spherules_near_the_Kamil_crater

Cronin, J. R., Pizzarello, S. and Cruikshank, D. P. (1988) Organic matter in carbonaceous chondrites, planetary satellites, asteroids, and comets. pp. 819–857 of JF Kerridge and MS Matthews, Editors, Meteorites and the Early Solar System, Univ. Arizona Press.

de Bosnia, P. (2017, agosto 20). MISTERIOS DEL VALLE DE LA MUERTE – Parte 4 (última). PIRAMIDES DE BOSNIA. https://piramidesdebosnia.com/2017/08/20/misterios-del-valle-de-la-muerte-parte-4-ultima/

de Vera, JP. (2015). Panspermia. In:, et al. Encyclopedia of Astrobiology. Springer, Berlin, Heidelberg. https://doi-org.ezproxy.utp.edu.co/10.1007/978-3-662-44185-5_1151

Di Rienzo J.A., Casanoves F., Balzarini M.G., Gonzalez L., Tablada M., Robledo C.W. InfoStat versión 2019. Centro de Transferencia InfoStat, FCA, Universidad Nacional de Córdoba, Argentina. URL http://www.infostat.com.ar

Griffin, D.W., 2004, Terrestrial microorganisms at an altitude of 20,000m in Earth’s atmosphere. Aerobiologia, 20, 135–140.

Griffin,D.W., 2008, None-spore forming eubacteria isolated at an altitude of 20,000m in Earth’s atmosphere: extended incubation periods needed for culture based assays. Aerobiologia, 24, 19–25.

Harris, M. J., Wickramasinghe, N. C., Lloyd, D., et al., 2002, Detection of living cells in stratospheric samples. Proceedings of SPIE Conference, 4495, 192–198.

Hoover R., Jerman, G., Rossignold-Strick, M. (2005). The Hollow Spheres of the Orgueil Meteorite: A Re-Examination. Proceedings of SPIE - The International Society for Optical Engineering. 5906. 10.1117/12.624859.

Hoyle, F., Wickramasinghe, C. (1981). Comets - A Vehicle for Panspermia. In: Ponnamperuma, C. (eds) Comets and the Origin of Life. Proceedings of the College Park Colloquia, vol 5. Springer, Dordrecht. 10.1007/978-94-009-8528-5_15

Imshenetsky A., Lysenko S., Kazakov G., 1978. Upper boundary of the biosphere. Appl.Environ.Microbiol., 35, 1–5.

Irvine, W.M. (2015). Astrobiology. In:, et al. Encyclopedia of Astrobiology. Springer, Berlin, Heidelberg. https://doi-org.ezproxy.utp.edu.co/10.1007/978-3-662-44185-5_120.

Joseph, R., Impey, C., Planchon, O., del Gaudio, R., Abu Safa, M., Sumanarathna, A. R., Ansbro, E., Duvall, D., Bianciardi, G., Gibson, C. H., & Schild, R. (2024). Extraterrestrial Life in the Thermosphere: Plasmas, UAP, Pre-Life, Fourth State of Matter. *Journal of Modern Physics*, *15*, 322–374.

Kos, S., Zupančič, N., Gosar, M., & Miler, M. (2022). Solid Carriers of Potentially Toxic Elements and Their Fate in Stream Sediments in the Area Affected by Iron Ore Mining and Processing. Minerals, 12(11), 1424.

Rossignol-Strick, M. and Barghoorn E.S. (1971) Extraterrestrial abiotic organization of organic matter: the hollowspheres of the Orgueil meteorite, Space Life Sci., 3, 89–107.

Schmitt-Kopplin, P., et al, (2010) High molecular diversity of extraterrestrial organic matter in Murchison meteorite revealed 40 years after its fall, Proc.Nat.AcadSci., 107, No.7. 2763–2768

Sergievskii V. (2020) Super Strong Artificial Intelligence and Human Mind, Procedia Computer Science, Volume 169, Pages 458-460, 10.1016/j.procs.2020.02.225.

(s/f). Academia Rusa de Ciencias. Polvo cósmico en los ventisqueros de la Antártida. Recuperado el 13 de junio de 2024, de https://new.ras.ru/activities/news/kosmicheskaya-pyl-v-sugrobakh-antarktidy/

(S/f). Eleconomista.net. Recuperado el 13 de junio de 2024, de https://www.eleconomista.net/deportes/Meteoritos-de-2700-millones-de-anos-revelan-secretos-de-la-antigua-atmosfera-20160511-0022.html

Shivaji, S., Chaturvedi, P., Begum, Z. et al, 2009. Janibacter hoylei sp. nov., Bacillus isronensis sp. nov. and Bacillus aryabhattai sp. nov., isolated from cryotubes used for collecting air from the upper atmosphere, Int. J. System.Evolut.Microbiol., 59, 2977–2986.

Smith, D.J., Griffin, D.W & Schuerger A.C., 2010. Stratospheric microbiology at 20km over the Pacific Ocean. Aerobiologia, 26, 35–46.

Staley J. 2003. Astrobiology, the transcendent science: the promise of astrobiology as an integrative approach for science and engineering education and research, Current Opinion in Biotechnology, Volume 14, Issue 3, Pages 347-354, 10.1016/S0958-1669(03)00073-9.

Suttle, M., Campanale, F., Folco, L., Tavazzani, L., Meier, M., Miller, C., Hughes, G., Genge, M., Salge, T., Spratt, J., & Anand, M. (2023). Fossil micrometeorites from Monte dei Corvi: Searching for dust from the Veritas asteroid family and the utility of micrometeorites as a palaeoclimate proxy. Geochimica Et Cosmochimica Acta, 355, 75–88. 10.1016/j.gca.2023.06.027

Wainwright M. (2015) Biological entities and DNA-containing masses isolated from the stratosphere-evidence for a non-terrestrial origin, Astronomical Review, 11:2, 25–40, DOI: 10.1080/21672857.2015.1087751

Wainwright M., Rose C., Baker A., Wickramasinghe C. 2013a. More biological entities from the stratosphere including a diatom fragment-further evidence for a space origin. Journal of Cosmology, Vol.,22, pp 10267-10274.

Wainwright M., Rose C., Baker A., Wickramasinghe C. 2013b. Microspherules and Preumptive Biological Entities Found Inside the Polonnaruwa Meteorite. Journal of Cosmology, Vol.,22, pp 10197-10202

Wainwright M., Rose C., Baker A., Wickramasinghe C. 2013c. Biological Entities Isolated from the Stratosphere (22-27km) –Case for their Space Origin.

Instruments, Methods, and Missions for Astrobiology XVI, edited by Richard B. Hoover, Gilbert V. Levin, Alexei Yu. Rozanov, Nalin C. Wickramasinghe, Proc. of SPIE Vol. 8865, 88650L doi: 10.1117/12.2047304

Wainwright M., Rose C., Baker A., Wickramasinghe C. 2013d. Isolation of biological entities from the stratosphere (22-27Km) Journal of Cosmology, Vol.,22, pp 10189-10197

Wainwright M., Rose C., Baker A., Wickramasinghe C. 2013e Isolation Of A Diatom Frustule Fragment From The Lower Stratosphere (22–27km)-Evidence For A Cosmic Origin. Journal of Cosmology, 22, pp. 10183–10186, DOI: 10.5334/jc.sf

Wainwright M., Rose C., Baker A., Wickramasinghe C. 2013f Filamentous Biological Entities Obtained from the Stratosphere Journal of Cosmology, Vol.,22, pp 10206-10210.

Wainwright M., Rose C., Baker A., Wickramasinghe C. 2013g Biology Associated With A Titanium Sphere Isolated From The Stratosphere. Journal of Cosmology 23, pp.1117–11124.

Wainwright M., Rose C., Baker A., Briston K., Wickramasinghe C. (2013h) Allen Hills and Schopf-like putative fossilized bacteria seen in a new type of carbonaceous meteorite. Journal of Cosmology, 22, pp. 10198–10201, DOI: 10.5334/jc.sh

Wainwright M., Wickramasinghe C. (2021) Polonnaruwa Stones Revisited – Evidence for Non-Terrestrial Life Advances in Astrophysics, Vol. 6, No. 2, 10.22606/adap.2021.62001

Wainwright M., Wickramasinghe C., Rose C., Baker A. (2014) Recovery of Cometary Microorganisms from the Stratosphere. Astrobiol Outreach 2:110. doi: 10.4172/2332-2519.1000110

Wainwright M., Wickramasinghe C., Tokoro G. (2021) Neopanspermia – Evidence That Life Continuously Arrives at the Earth from Space Advances in Astrophysics, Vol. 6, No. 1, February 2021 10.22606/adap.2021.61002Advanc

Wainwright, M., Wickramasinghe, N.C., Narlikar, J.V. & Rajaratnam, P.,2003. Microorganisms cultured from stratospheric air samples obtained at 41 km. FEMS Microbiol Lett, 218, 161- 165.Ward, J.H. (1963). Hierarchical Grouping to Optimize an Objective Function. Journal of the American Statistical Association, 58: 236–244

Wesson, P.S. Panspermia, Past and Present: Astrophysical and Biophysical Conditions for the Dissemination of Life in Space. Space Sci Rev 156, 239–252 (2010). https://doi-org.ezproxy.utp.edu.co/10.1007/s11214-010-9671-x

Wickramasinghe C., Tokoro G., Wainwright M. (2015) The Transition from Earth-centred Biology to Cosmic Life #. JAstrobiolOutreach 3: 122. doi:10.41721/2332-2519.1000122

Yamagishi, A. (2019). What Is Astrobiology?. In: Yamagishi, A., Kakegawa, T., Usui, T. (eds) Astrobiology. Springer, Singapore. https://doi-org.ezproxy.utp.edu.co/10.1007/978-981-13-3639-3_1

Yang, Y., Itahashi, S., Yokobori, S. & Yamagishi, A., 2008. UV-resistant bacteria isolated from upper troposphere and lower stratosphere. Biol. Sci. in Space, 22, 18–25.

Yang, Y., Yokobori, S. & Yamagishi, A., 2009. Assessing panspermia hypothesis by microorganisms collected from the high altitude atmosphere, Biol. Sci. in Space, 23, 151–163.

